# Fibronectin-Assisted Coating of Low-Molecular-Weight Fucoidan Creates a Biomimetic Interface for Endothelialization of Cobalt-Chromium Stents

**DOI:** 10.64898/2026.11.02.763584

**Authors:** Mehdi Saadi, Raymond Haddad, Nour Zeaiter, Oualid Haddad, Divya Prabhakar Reddy, Jean Sébastien Baumann, Fahima Irki, Jessica Bamba-Funck, Angela Sutton, Céline Falentin-Daudré, Kévin Bassand, Catherine Boisson Vidal, Hanna Hlawaty

## Abstract

Incomplete or delayed endothelialization after stent implantation compromises vascular healing and increases the risk of restenosis and late thrombosis. To address this limitation, Cobalt-Chromium (Co-Cr) stents were modified with a biomimetic coating combining Low-Molecular-Weight Fucoidan (LMWF), a sulfated polysaccharide with glycocalyx-mimetic properties, and fibronectin, to promote endothelial repair while preserving vascular homeostasis. The morphology, chemical composition, wettability and stability of the coating were characterized using complementary physicochemical approaches. The behavior of human endothelial cells cultured on this dual fibronectin-LMWF-coated stent was then assessed.

Our results showed that fibronectin-LMWF-based coating achieved a ten-fold higher surface density on the stent than LMWF alone (0.29 ± 0.04 *vs* 0.03 ± 0.01 µg/mm^2^) and retained 0.27 ± 0.03 µg/mm^2^ of the coating after 168 h, compared with 0.01 ± 0.01 µg/mm^2^ for LMWF alone.

Although wettability remained similar to that of uncoated stents, the combined coating markedly enhanced endothelial cell adhesion, spreading, lamellipodia formation, proliferation, and MMP-2 secretion. These effects were accompanied by enhanced F-actin organization and redistribution of membrane-associated VE-cadherin, integrins β1 and α6, and syndecans-1 and-4, without major changes in their total expression. Importantly, the coating induced these responses without detectable cytotoxicity or significant inflammatory activation.

Overall, the fibronectin-LMWF-based coating provides a stable biomimetic interface that enhances endothelialization through coordinated remodeling of adhesion, cytoskeletal, and glycocalyx-associated structures, while maintaining endothelial homeostasis on Co-Cr stent. This strategy offers a promising platform for next-generation glycocalyx-like stent coatings designed to improve vascular healing and limit restenosis.

**Graphical abstract:** Fibronectin-assisted immobilization of low-molecular-weight fucoidan enhances endothelialization of Cobalt-Chromium stents. Schematic illustration of the sequential dip-coating modification of Cobalt-Chromium (Co-Cr) stents with fibronectin and low-molecular-weight fucoidan (LMWF), followed by HUVECs seeding and endothelial colonization. Fibronectin-assisted LMWF immobilization enhances coating retention and supports HUVECs adhesion, spreading, and proliferation. Representative microscopy images illustrate coating retention, endothelial morphology, surface colonization, and VE-cadherin distribution. The proposed coating architecture involves fibronectin adsorption onto the Co-Cr surface, followed by the formation of electrostatic interactions between fibronectin and LMWF as well as between LMWF and endothelial cell-surface components (selectins). Overall, these findings highlight the potential of fibronectin-LMWF coatings as a biomimetic strategy to promote Co-Cr stent endothelialization. Created in BioRender. Saadi, M. (2027) https://BioRender.com/6xb50t5.

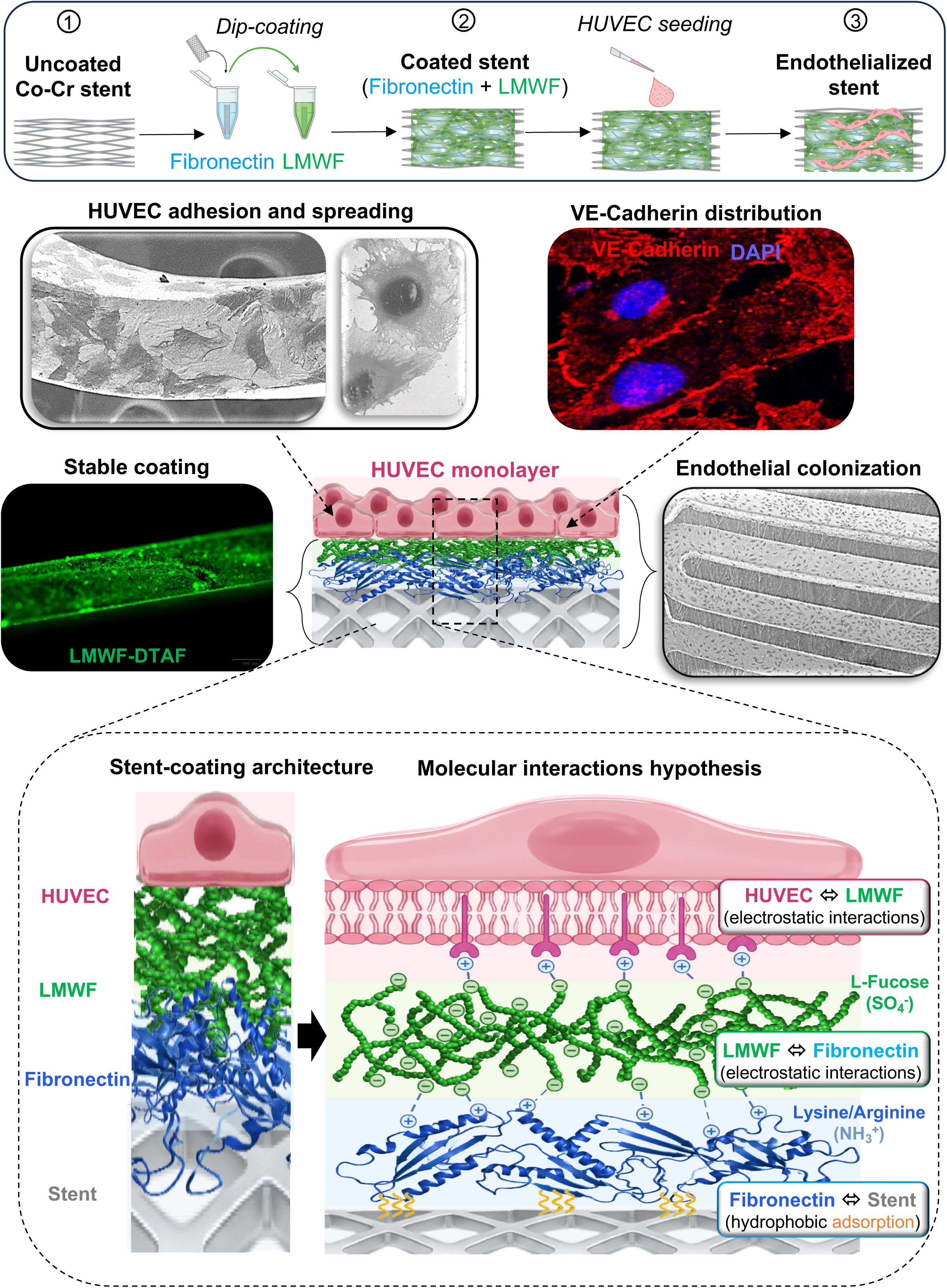

## 1. Introduction

Stent implantation is an established minimally-invasive treatment for vascular stenosis. However, vascular injury as well as delayed or incomplete endothelial healing following stent implantation can promote inflammation, neointimal hyperplasia, and thrombotic complications [1–4]. In-stent restenosis remains a major limitation to the long-term clinical outcomes, primarily driven by vascular smooth muscle cell (SMC) migration into the intima and subsequent proliferation, leading to luminal narrowing [4–6]. Endothelial injury also promotes platelet activation and leukocyte recruitment, further amplifying the inflammatory response and neointimal growth [2,7,8]. Although drug-eluting stents (DES) reduce restenosis by locally releasing antiproliferative agents, these drugs can also delay endothelial repair and vascular healing [1,9]. This therapeutic trade-off has motivated the development of new-generation bioactive stent coatings designed to accelerate endothelialization while limiting neointimal growth [10,11].

Fucoidans are sulfated polysaccharides derived from brown seaweeds [12]. Among them, Low-Molecular-Weight Fucoidan (LMWF) has emerged as a promising candidate for vascular repair owing to its ability to modulate endothelial and SMC behavior [13]. We previously demonstrated that, *in vitro*, LMWF promotes endothelial cell migration while inhibiting SMC migration, effects associated with increased matrix metalloproteinase-2 (MMP-2) expression in endothelial cells and decreased expression in SMCs [14]. LMWF has also been shown to enhance endothelial progenitor cell (EPC) proliferation and migration and to promote vascular cord formation in the presence of fibroblast growth factor 2 (FGF-2), in association with increased integrin α6 expression [15]. Furthermore, our previous work showed that LMWF-induced endothelial migration and angiogenesis involve heparanase and syndecan-4 (SDC-4), implicating extracellular matrix (ECM) remodeling and adhesion-associated pathways in these processes [16]. Consistent with its potential to modulate vascular remodeling, we also demonstrated that LMWF reduced neointimal hyperplasia in a rat model of arterial injury [14].

Beyond its biological effects, fucoidan has been investigated as a component of bioactive vascular coatings. Its incorporation into vascular endothelial growth factor (VEGF)-based polyelectrolyte multilayer coatings on decellularized porcine pulmonary valves improved endothelial cell adhesion and colonization [17]. Similarly, dextran-graft-polybutylmethacrylate (Dex-PBMA) copolymer coatings on Cobalt-Chromium (Co-Cr) stents showed promising *in vitro* and *in vivo* performance, including improved re-endothelialization and preservation of the vessel lumen [18]. More recently, Bai *et al.* developed a fucoidan-polydopamine-based coating on a Mg-Zn-Y-Nd alloy, reporting improved corrosion resistance, hemocompatibility, and endothelialization, together with reduced intimal hyperplasia and tissue inflammation [19]. Collectively, these findings highlight the potential of polysaccharides as bioactive coating components for vascular applications and support further investigation of LMWF-based surface modification to promote endothelialization of metallic vascular scaffolds.

However, translating LMWF-based functionalization to metallic stents requires effective surface deposition and sustained coating retention [2]. Combining LMWF with an adhesive ECM protein such as fibronectin may help achieve these objectives while promoting endothelial cell attachment. Fibronectin promotes endothelial adhesion through integrin-mediated interactions and contains a Heparin-binding domain II (Hep-II) that binds heparan sulfate [20]. Given the structural and functional similarities between fucoidan and sulfated glycosaminoglycans (GAGs), including their interactions with certain heparin-binding proteins, fibronectin may provide a favorable interfacial layer for LMWF retention while supporting endothelial colonization. However, whether such interactions contribute to coating stability remains to be established. Characterizing the physicochemical properties of functionalized surfaces is essential for relating coating composition to biological performance. Fourier-transform infrared spectroscopy (FT-IR) identifies characteristic chemical groups associated with coating components [21], whereas energy-dispersive X-ray spectroscopy (EDS) provides information on surface elemental composition [22]. Water contact angle measurements further assess surface wettability, helping distinguish hydrophilic from hydrophobic surface characteristics [21]. Together, these complementary techniques provide a physicochemical profile of the coated surfaces before biological evaluation.

We hypothesized that fibronectin precoating would enhance LMWF retention on Co-Cr stents and promote endothelial colonization. To test this hypothesis, we developed a sequential fibronectin-LMWF coating and characterized its physicochemical properties and stability. We then assessed endothelial cell adhesion, spreading, proliferation, lamellipodia formation, cytoskeletal organization, and adhesion-associated protein distribution, alongside cytotoxicity and inflammatory responses. Finally, we investigated whether conditioned media from endothelial cells modulated SMC migration and metabolic activity.

## 2. Materials and methods

### 2.1. Cell Culture

#### 2.1.1. Endothelial cell culture

Human umbilical vein endothelial cells (HUVECs; C-12200, PromoCell GmbH, Germany) were cultured in Endothelial Cell Basal Medium 2 (ECBM-2; C-22111, PromoCell GmbH, Germany) supplemented with 12 % fetal bovine serum (FBS), recombinant human epidermal growth factor (rhEGF, 5 ng/mL), recombinant human basic fibroblast growth factor (rh-FGF-2, 10 ng/mL), recombinant human insulin-like growth factor (20 ng/mL), recombinant human vascular endothelial growth factor 165 (0.5 ng/mL), ascorbic acid (1 µg/mL), hydrocortisone (0.2 µg/mL) and heparin (22.5 µg/mL).

#### 2.1.2. Smooth muscle cell culture

Human smooth muscle cells (HSMCs; C-12533, PromoCell GmbH, Germany) were cultured in Smooth Muscle Cell Growth Medium 2 (SMCsG-2; C-22162, PromoCell GmbH, Germany) supplemented with 5 % FBS, rhEGF (0.5 ng/mL), rh-FGF-2 (2 ng/mL), and recombinant human insulin (5 µg/mL). Cells were maintained under standard culture conditions (37 °C, 5 % CO_2_).

### 2.2. Cobalt-Chromium stents

The Optimus^®^ Cobalt-Chromium (Co-Cr) stent (L version, 9 Zigs) was provided by AndraTec Medical Devices GmbH (Koblenz, Germany). Each stent was cut into eight segments, each measuring 5 × 0.5 × 0.22 mm, with a nominal surface area of 7.2 mm² per segment, accounting for all four faces. These stents had previously been used in interventions for congenital heart disease [23,24,25].

### 2.3. Stent coating

#### 2.3.1. Coating components: fibronectin and fucoidan

Human fibronectin (440,000 g/mol; Corning Incorporated, United States). Low-Molecular-Weight Fucoidan (LMWF; 15,500 g/mol) and 5-([4,6-dichlorotriazin 2-yl]-amino) fluorescein (DTAF)-labelled LMWF (LMWF-DTAF; 15,500 ± 3,000 g/mol) were kindly provided by Dr C. Boisson-Vidal (UMRS-1144, Paris, France) [12,26,27]. Dextran (550,000 g/mol; Amersham, United Kingdom) was used as a coating control.

#### 2.3.2. Dip-Coating methods

Coating solutions were prepared in phosphate-buffered saline (PBS, pH 7.4). Stents were coated by dip coating using 100 µg/mL solutions of fibronectin or LMWF, either individually or sequentially to generate a bilayer fibronectin-LMWF coating. In the sequential condition, stents were first incubated with fibronectin and subsequently coated with LMWF (graphical abstract). Each coating step consisted of successive 10 sec incubations in 300 µL of coating solution. No rinsing was performed between incubations or after the final coating step. Coated stents were air-dried under sterile conditions before use.

For fluorescence imaging, fluorescently labelled LMWF-DTAF was applied either alone or after fibronectin precoating using the same procedure. Uncoated stents and dextran-coated stents served as controls.

#### 2.3.3. Coating retention

Coating retention was evaluated by monitoring the release of fluorescently labelled fucoidan (LMWF-DTAF) into PBS. Individual stents fragments were immersed in 200 µL of PBS in sterile 96-well plates. The fluorescence intensity of the release medium was measured using a TriStar^2^ LB 942 fluorometer (Berthold Technologies, Germany) at excitation and emission wavelengths of 495 and 521 nm, respectively. Fluorescence measurements were corrected for PBS background. Samples were maintained in PBS at 4 °C throughout the 7-days (168 h) retention experiment. They were kept under static conditions for the first 72 h, followed by orbital agitation at 23 rpm until 168 h, to promote medium mixing and assess the release of coating-associated components, following previously reported *in vitro* drug-release protocols for drug-eluting stents [28]. At each sampling time, the release medium was collected and replaced with fresh PBS.

A calibration curve (0-100 µg/mL LMWF-DTAF) was used to quantify fucoidan concentrations in the coating solutions and in the release medium. The deposited amount was determined from the decrease in the coating solution concentration, accounting for solution volumes, and normalized to the sample surface area (7.2 mm^2^ x number of segments) to obtain the surface density (µg/mm^2^). At each time point, the released amount was calculated from the measured concentration and the collected PBS volume. The remaining amount was obtained by subtracting cumulative release from the initial deposited amount, with all sampling time points included in the retention calculations. The remaining surface density was normalized to the sample surface area, and retention was expressed as a percentage of the initial deposited amount after PBS background correction. Only selected time points are shown in Figure 1C. Four independent samples were analyzed per condition (n = 4).

**Figure 1.**
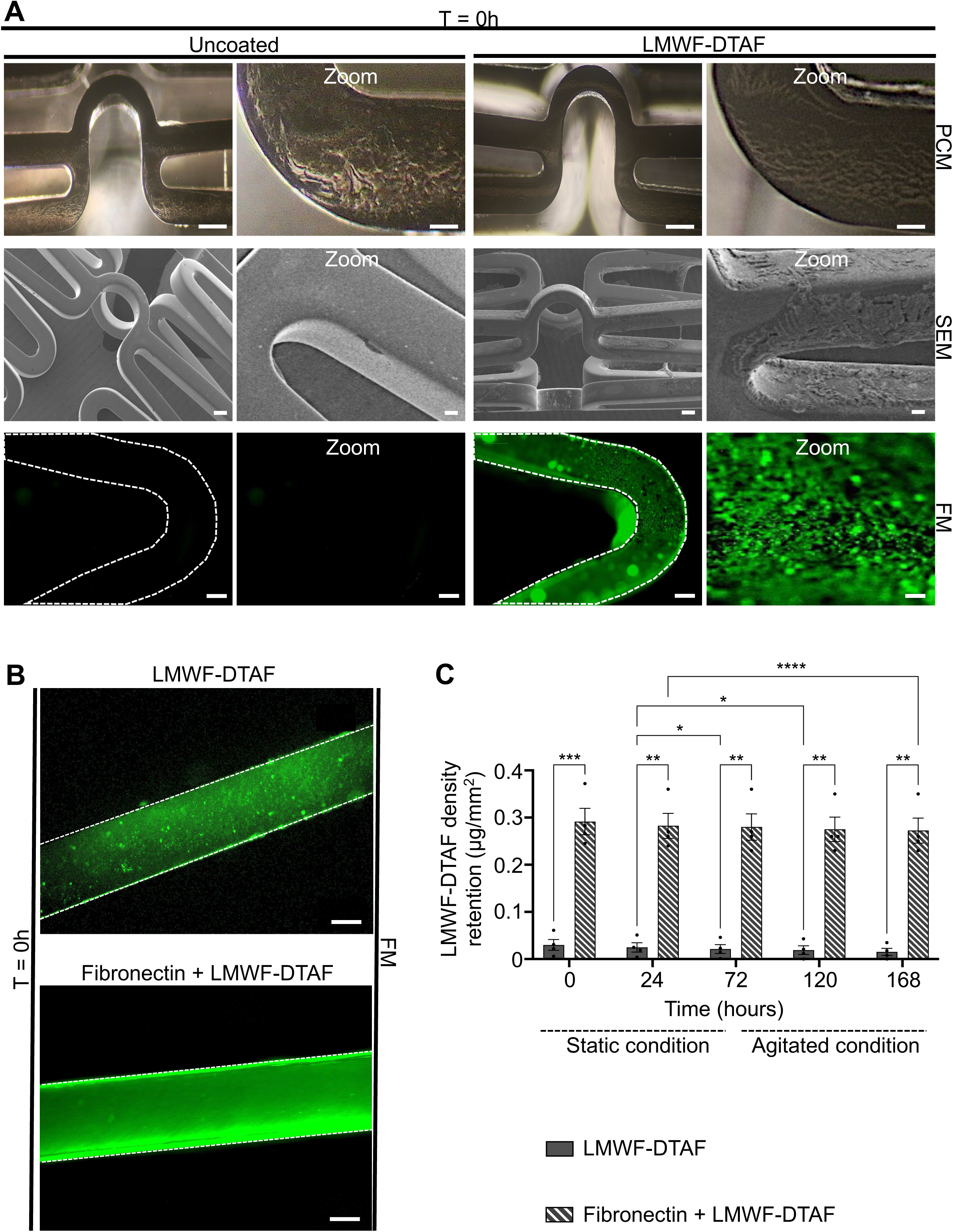
Fibronectin-assisted LMWF deposition and retention on Co-Cr stents. **(A)** Representative phase-contrast microscopy (PCM), scanning electron microscopy (SEM), and fluorescence microscopy (FM) images of uncoated and LMWF-DTAF-coated Co-Cr stents at T = 0h. PCM and SEM images show surface morphology of uncoated and coated stents, while FM images show the presence and surface distribution of fluorescently labeled LMWF-DTAF. Higher-magnification views are presented on the right. White dashed lines delineate the stent boundaries. Scale bars: PCM = 200 µm (overview) and 46 µm (higher-magnification); SEM = 200 µm (overview) and 40 µm (higher-magnification); FM = 100 µm (overview), and 10 µm (higher-magnification). **(B)** Representative FM images of LMWF-DTAF-coated and fibronectin-LMWF-DTAF-coated Co-Cr stents at T = 0 h. White dashed lines delineate the stent boundaries. Scale bars: 100 µm. **(C)** Quantification of LMWF-DTAF surface density retained on stents coated with LMWF-DTAF alone or sequentially with fibronectin followed by LMWF-DTAF over 168 h. Stents were maintained under static conditions from 0 to 72 h and under agitated conditions from 72 to 168 h. Data are mean ± SEM (n = 4). Two-way ANOVA with Tukey’s multiple-comparisons test. \**p* < 0.05, \*\**p* < 0.01, \*\*\**p* < 0.005, \*\*\*\**p* < 0.001.

#### 2.3.4. Physicochemical characterization

The chemical composition of the coatings was characterized by attenuated total reflectance Fourier-transform infrared spectroscopy (ATR-FTIR; Thermo Nicolet Avatar 370 spectrometer, Thermo Fisher Scientific, United States). Spectra were acquired directly from coated stent fragments. Uncoated fragments were analyzed under the same conditions as controls. The acquisition range was 4000 - 600 cm^-1^ with 64 scans and a spectral resolution of 4 cm^-1^. Sulfate-associated bands (1250 - 1100 cm^-1^) were examined to assess the presence of LMWF [29]. Amide I and amide II bands (1700 - 1600 cm^-1^ and 1580 - 1480 cm^-1^, respectively) were used to confirm the presence of fibronectin [30].

Surface morphology and coating homogeneity were examined by scanning electron microscopy (SEM; TM3000, Hitachi High-Technologies Corporation, Tokyo, Japan). Elemental composition and surface distribution were assessed by energy-dispersive X-ray spectroscopy (EDS) using a SwiftED3000 system (Oxford Instruments, Abingdon, United-Kingdom). Three independent stent fragments were analyzed per condition. A total of 23 - 60 EDS measurements were collected per condition across these fragments. Measurements were averaged within each fragment before statistical analysis (n = 3 fragments per condition, 23 - 60 EDS measurements each).

#### 2.3.5. Coating distribution

Coating distribution was evaluated using the ZOE^TM^ Fluorescent Cell Imager (Bio-Rad, United States). Stents were coated with LMWF-DTAF either alone or sequentially after fibronectin precoating. Fluorescence images were acquired to assess the distribution and apparent homogeneity of LMWF-DTAF over the stent surface.

#### 2.3.6. Hydrophilicity characterization

Surface hydrophilicity was evaluated by static water contact angle measurements using the sessile drop method and a DSA10 drop shape analyzer (KRÜSS, France). Ultrapure water droplets (0.2 µL) were deposited onto each stent surface. Three droplets were analyzed per sample, and the mean contact angle was calculated. Three independent samples were analyzed per condition (n = 3).

#### 2.3.7. pH stability

The effect of the different coatings on the pH stability of the culture medium was evaluated over 72 h in the absence of cells. Co-Cr stents were placed individually in sterile 24-well plates containing 4 mL of complete culture medium per well. A medium-only condition was included as a control. Plates were incubated at 37 °C in a humidified atmosphere containing 5% CO2. Each condition was tested in triplicate. The pH was measured directly in each well at 0.5, 24, 48 and 72 h using a FiveEasy benchtop pH meter (Mettler Toledo).

### 2.4. Stent endothelialization

#### 2.4.1. Endothelial cell seeding

Preliminary seeding experiments were performed on uncoated stents using three initial cell numbers: 1.5 x10^5^, 3.75 x10^5^ and 15 x10^5^ HUVECs per fragment (7.2 mm^2^/segment). Cell colonization was evaluated at 24 and 72 h to establish suitable seeding conditions (Fig. 3C-D). Based on these experiments, 3.75 x10^5^ cells per fragment were selected for subsequent assays. Uncoated, fibronectin-coated, LMWF-coated and fibronectin-LMWF-coated Co-Cr stent fragments were individually placed in 24-well culture plates. For fluorescence-based analyses of cell colonization, spreading, and lamellipodia formation, HUVECs were labelled with CellTracker^TM^ Green CMFDA (C7025, Thermo Fisher Scientific, United States; 1:2000 dilution). Labelling was performed in serum-free medium for 45 min at 37 °C before seeding. Cells were seeded onto each stent fragment in a 50 µL suspension and allowed to adhere for 4 h at 37 °C in a humidified atmosphere containing 5% CO_2_. Complete culture medium was then added to a final volume of 1 mL per well.

#### 2.4.2. Scanning electron microscope analysis

HUVECs-seeded Co-Cr stents were rinsed with PBS and fixed overnight at 4 °C in 2 % glutaraldehyde prepared in 0.1 M phosphate buffer (pH 7.4). Samples were washed with 0.1 M cacodylate buffer and post-fixed for 20 min in 1 % osmium tetroxide prepared in the same buffer. Dehydration was performed through a graded ethanol series (50 %, 70 %, 96 % and 100 %), followed by successive incubations in a 1:1 (v/v) ethanol/hexamethyldisilazane (HMDS) solution and pure HMDS. Samples were dried under vacuum in a desiccator, mounted on scanning electron microscope (SEM) stubs, and sputter-coated with platinum or tungsten (ACE600, Leica Microsystems). SEM observations were performed using a GeminiSEM 500 field-emission SEM (ZEISS) operated at 3 kV. Images were acquired using the InLens detector or the SE2 secondary electron detector.

#### 2.4.3. Cell colonization and proliferation

Initial cell attachment was evaluated at 4 h after seeding. Subsequent colonization was assessed at 24, 48 and 72 h by fluorescence microscopy (FM). Cell density was evaluated in four independent experiments (n = 4), each including three stent fragments per condition as technical replicates. Five fields were analyzed per fragment. In each field, HUVECs were counted and the imaged stent surface area was measured manually using ImageJ software (version 1.54p, National Institutes of Health, Bethesda, MD, United States). The density for each fragment was calculated by dividing the total cell count across the five fields by their combined surface area. The three replicate densities were then averaged to obtain one value per independent experiment. Cell density was expressed as cell/mm^2^.

Proliferative activity was assessed by 5-ethynyl-2′-deoxyuridine (EdU) incorporation using the Click-iT^TM^ EdU Flow Cytometry Assay Kit (C10418, Thermo Fisher Scientific, United States), adapted for FM. For this assay, HUVECs were synchronized by overnight serum starvation and seeded as described above. After 72 h, the medium was replaced with fresh medium containing 10 µM EdU. After a further 24 h, samples were washed with PBS and fixed with 4% paraformaldehyde (PFA) for 10 min at room temperature (RT). They were then permeabilized with 0.01% Triton X-100 for 5 min and washed again with PBS. Fluorescence images were acquired using the ZOE^TM^ Fluorescent Cell Imager (Bio-Rad, United States). For the EdU assay, three independent stents were analyzed per condition. Five randomly selected fields were examined per sample. Proliferative activity was expressed as the percentage of EdU-positive cells among the total number of HUVECs.

#### 2.4.4. Cytotoxicity assay

Cytotoxicity was evaluated by measuring lactate dehydrogenase (LDH) release into serum-free conditioned medium collected after 24, 48 and 72 h of culture. LDH activity was quantified using the CytoTox 96^®^ Non-Radioactive Cytotoxicity Assay (Promega, G1780, United States) according to the manufacturer’s instructions. Briefly, 50 µL of each supernatant was analyzed in triplicate in a 96-well plate. Absorbance was measured at 490 nm using a microplate reader. Maximum LDH release was determined from lysed HUVECs and used as the positive control. Three independent biological replicates were analyzed per condition (n = 3).

#### 2.4.5. Cell spreading, lamellipodia formation and cytoskeletal organization

Cell spreading and lamellipodia formation were evaluated on uncoated and coated Co-Cr stents after 4 and 72 h of culture. HUVECs were labelled with CellTracker^TM^ Green CMFDA as described above. FM images of the labelled cells were used for quantitative analyses. Cell spreading was quantified by tracing individual cell outlines and measuring their projected area using ImageJ. Eighty cells were analyzed per image, with five randomly selected fields examined per stent. Cell area was expressed in µm^2^. Lamellipodia were quantified as the number of protrusions per field using the same fluorescence images. Four independent stents were analyzed per condition (n = 4). Representative SEM images were acquired at 48 h to visualize cell morphology and membrane protrusions. F-actin organization was examined after fixation and permeabilization by staining with Alexa Fluor^TM^ 568-conjugated phalloidin (1:400; A12380, Thermo Fisher Scientific, United States). Nuclei were counterstained with DAPI (10 µg/mL, 15 min, 37 °C, in the dark).

#### 2.4.6. Immunocytochemistry

The distribution of proteins involved in endothelial cell-cell and cell-matrix interactions was analyzed by immunocytochemistry after 48 h of culture. Cell-cell interactions were assessed by immunostaining for VE-cadherin (sc9989, Santa Cruz Biotechnology, United States). Cell-matrix interactions were assessed using antibodies against integrin β1 (sc374429), integrin α6 (sc374057), syndecan-1 (SDC-1; sc12765), and syndecan-4 (SDC-4; sc12766) (Santa Cruz Biotechnology, United States). Cells were fixed with 4 % PFA for 20 min at 4 °C, washed three times with PBS, and incubated for 1 h at RT in 0.2% gelatin to block non-specific binding. Samples were incubated overnight at 4 °C with primary antibodies (4 µg/mL) in a humidified chamber. Following three PBS washes, samples were incubated for 1 h at RT in the dark with Alexa Fluor^TM^ 555-conjugated goat anti-mouse IgG (4 µg/mL; Thermo Fisher Scientific, United States). Isotype-matched mouse IgG2a_k_ and rat IgG1_k_ antibodies were used as negative controls. Nuclei were counterstained with DAPI.

#### 2.4.7. Western blotting analysis

Whole-cell proteins were extracted after 48 h of culture using Pierce™ RIPA buffer (Ref. 89900, Thermo Fisher Scientific, United States) supplemented with protease and phosphatase inhibitors. Protein extracts were concentrated using Amicon^®^ Ultra-0.5 centrifugal filters with a 30 kDa molecular-weight cutoff (Merck, Germany). Protein concentration was determined using the Pierce^TM^ BCA Protein Assay Kit (Thermo Fisher Scientific, United States). Thirty µg of total protein were separated by SDS-PAGE on 10 % polyacrylamide gels at 100 V for 90 min and transferred onto 0.2 µm nitrocellulose membranes (Amersham Protran^TM^, Cytiva, United Kingdom) at 150 V for 1 h. Membranes were blocked for 1 h at RT in TBS-T containing 5% bovine serum albumin (BSA) and incubated overnight at 4 °C with primary antibodies.

Mouse monoclonal antibodies against VE-cadherin (sc9989), integrin β1 (sc374429) and integrin α6 (sc374057; all from Santa Cruz Biotechnology, United States) at a concentration of 0.4 µg/mL, and β-actin (sc8432) at a concentration of 0.2 µg/mL. Rabbit monoclonal antibodies against phospho-paxillin (Tyr118; E9U9F) and total paxillin (D9G12, Cell Signaling Technology, United States) were diluted 1:1000. After washing with TBS-T, membranes were incubated for 1 h with the corresponding HRP-conjugated goat anti-mouse or goat anti-rabbit secondary antibodies (DAKO Agilent, United States; 1:1000). Immunoreactive bands were detected using Pierce^TM^ ECL Western Blotting Substrate (Thermo Fisher Scientific, United States) and visualized with a ChemiDoc^TM^ imaging system (Bio-Rad, United States). When required, membranes were stripped using a RestoreTM Western Blotting Stripping Buffer (Thermo Fisher Scientific, United States). Band intensity was quantified using Image Lab software (Bio-Rad, United States).

#### 2.4.8. Dot Blot analysis

Conditioned medium collected after 48 h of culture was centrifuged at 300 x g for 5 min to remove cellular debris. Protein concentration was determined using the Pierce^TM^ BCA Protein Assay Kit. Equal amounts of protein (75 µg) were spotted directly onto Hybond^TM^-N+ nitrocellulose membranes (Amersham Protran^TM^, Cytiva, United Kingdom). Membranes were blocked for 1 h in PBS-T containing 0.5 % BSA and 3 % non-fat dry milk before overnight incubation at 4°C with anti-SDC-1 or anti-SDC-4 antibodies. Following PBS-T washes, membranes were incubated for 1 h with an HRP-conjugated goat anti-mouse secondary antibody (DAKO Agilent). HUVECs stimulated with 1 µM phorbol 12-myristate 13-acetate (PMA) served as a positive control for SDC shedding. Chemiluminescent signals were detected using an enhanced chemiluminescence substrate (Thermo Fisher Scientific, United States) and acquired with a ChemiDoc^TM^ imaging system (Bio-Rad, United States). Signal intensity was quantified using Image Lab software (v6.1; Bio-Rad). Six independent samples were analyzed per condition (n = 6).

#### 2.4.9. Secretome analysis

HUVECs were cultured for an initial 24 h in complete medium. The medium was then replaced with serum-free medium. Conditioned medium was collected after 24 and 48 h of serum-free culture, corresponding to 48 and 72 h of total culture time, respectively. Samples were stored at-80 °C until analysis. Multiplex quantification of growth factors (FGF-2 and VEGF-A), chemokines (MCP-1/CCL2, RANTES/CCL5, and IL-8/CXCL8), cytokines (IL-6, TNF-α, and IL-10), and matrix metalloproteinases (MMP-2 and MMP-9) was performed using the U-PLEX^®^ Immuno-Oncology Group 1 Human Assay (Meso Scale Discovery, United States), according to the manufacturer’s instructions. Electrochemiluminescence signals were acquired using a QuickPlex SQ 120 reader (Meso Scale Discovery), and analyte concentrations were determined from the corresponding standard curves using the manufacturer’s analysis software.

### 2.5. Biological evaluation of SMCs response

#### 2.5.1. Migration assay

SMC migration was evaluated using a Boyden chamber fitted with polycarbonate membranes containing 8 µm pores (Corning Incorporated, United States). Membranes were coated with fibronectin (100 µg/mL) for 1 h at 37 °C before cell seeding. SMCs were seeded into the upper chambers at 6 x10^4^ cells in 500 µL of complete SMC growth medium. The lower chamber contained 500 µL of conditioned medium collected from HUVECs cultured on uncoated, fibronectin, LMWF, and fibronectin-LMWF-coated Co-Cr stents. After 24 h of incubation, non-migrated cells were removed from the upper surface of the membrane using a cotton swab. Migrated cells were fixed, stained with DAPI and imaged using a ZOE^TM^ Fluorescent Cell Imager (Bio-Rad, United States). Five randomly selected fields were analyzed per insert using ImageJ. Migration was expressed as the number of migrated SMCs per field.

#### 2.5.2. Metabolic activity assay

SMC metabolic activity was evaluated using an MTT assay after exposure to conditioned media collected from HUVECs cultured on uncoated, fibronectin-coated, LMWF-coated and fibronectin-LMWF-coated Co-Cr stents. SMCs were seeded in 96-well plates at 5 x10^3^ cells per well and cultured for 24 h in complete SMC growth medium to allow cell attachment. The medium was then replaced with a 1:1 mixture HUVEC-conditioned medium and fresh SMC growth medium. After 24 h of exposure, 10 µL of MTT solution (5 mg/mL in PBS; Sigma-Aldrich, United States) was added to each well and incubated for 3 h at 37 °C. The medium was removed, and the formazan crystals were dissolved in 100 µL of dimethyl sulfoxide (DMSO). Absorbance was measured at 570 nm. Three independent biological replicates were performed, with each condition analyzed in technical triplicate.

### 2.6. Statistical tests

Statistical analyses were performed using GraphPad Prism 10. Data are presented as mean ± SEM, with the number of independent biological replicates indicated in the corresponding figure legends. Technical replicates were averaged before statistical analysis. One-way ANOVA followed by Tukey’s multiple-comparisons test was used for comparisons involving a single experimental factor. Two-way ANOVA followed by Tukey’s multiple-comparisons test was used to assess the effects of coating condition and time. For non-parametric comparisons, the Kruskal-Wallis test followed by Dunn’s multiple-comparisons test was used. A two-sided *p* value < 0.05 was considered statistically significant.

## 3. Results

### 3.1. Fibronectin pre-coating increases LMWF deposition and retention on stents

The coating morphology and Low-Molecular-Weight Fucoidan (LMWF) labelled with 5-([4,6-dichlorotriazin 2-yl]-amino) fluorescein (LMWF-DTAF) distribution were first examined by phase-contrast microscopy (PCM), scanning electron microscopy (SEM), and fluorescence microscopy (FM) (Fig. 1A-B). Uncoated stents exhibited the characteristic reflective metallic surface. In contrast, coated stents showed reduced brightness under PCM, and SEM revealed a coating layer on both flat and curved regions of the stent struts (Fig. 1A). FM revealed no detectable fluorescence on uncoated stents, whereas LMWF-DTAF fluorescence was detected on stents coated with LMWF-DTAF alone and on those pre-coated with fibronectin before LMWF-DTAF deposition (Fig. 1A-B). LMWF-DTAF surface density was quantified using a fluorescence plate reader immediately after coating and during subsequent incubation in PBS (Fig. 1C). At T=0 h, fibronectin precoating was associated with approximately ten-fold higher LMWF-DTAF surface density than LMWF-DTAF coating alone (0.29 ± 0.04 µg/mm^2^ *vs* 0.03 ± 0.01 µg/mm^2^, \*\*\**p* < 0.005).

Furthermore, surface density was monitored at several time points up to 168 h, with or without agitation, and showed higher LMWF-DTAF surface density with fibronectin-precoating at all analyzed time points. At 24 and 72 h, surface density was 0.28 ± 0.03 and 0.28 ± 0.04 µg/mm^2^ respectively, after fibronectin precoating, compared with 0.02 ± 0.01 and 0.02 ± 0.01 µg/mm^2^ after LMWF-DTAF coating alone (\*\**p* < 0.01). Under agitation (from 72 h to 168 h), the corresponding-surface density values at 120 and 168 h were 0.27 ± 0.03 for both after fibronectin precoating, compared with 0.01 ± 0.01 for both after LMWF-DTAF coating alone (\*\**p* < 0.01). Overall, fibronectin-precoated stents retained more LMWF-DTAF per unit surface area than stents coated with LMWF-DTAF alone throughout the 168 h experiment.

### 3.2. Physicochemical characterization of coated stents

The chemical and elemental compositions, as well as the wettability of the coated surfaces, were characterized (Fig. 2A–C). Non-sulfated, electrically neutral dextran was included as a control to distinguish the effects of polysaccharide coating from those specifically associated with the negatively charged sulfate groups of LMWF, thereby assessing the contribution of sulfation to surface modification and wettability (hydrophilicity). The effects of the different coatings on culture medium pH were also evaluated under cell-free conditions (Fig. 2D).

**Figure 2.**
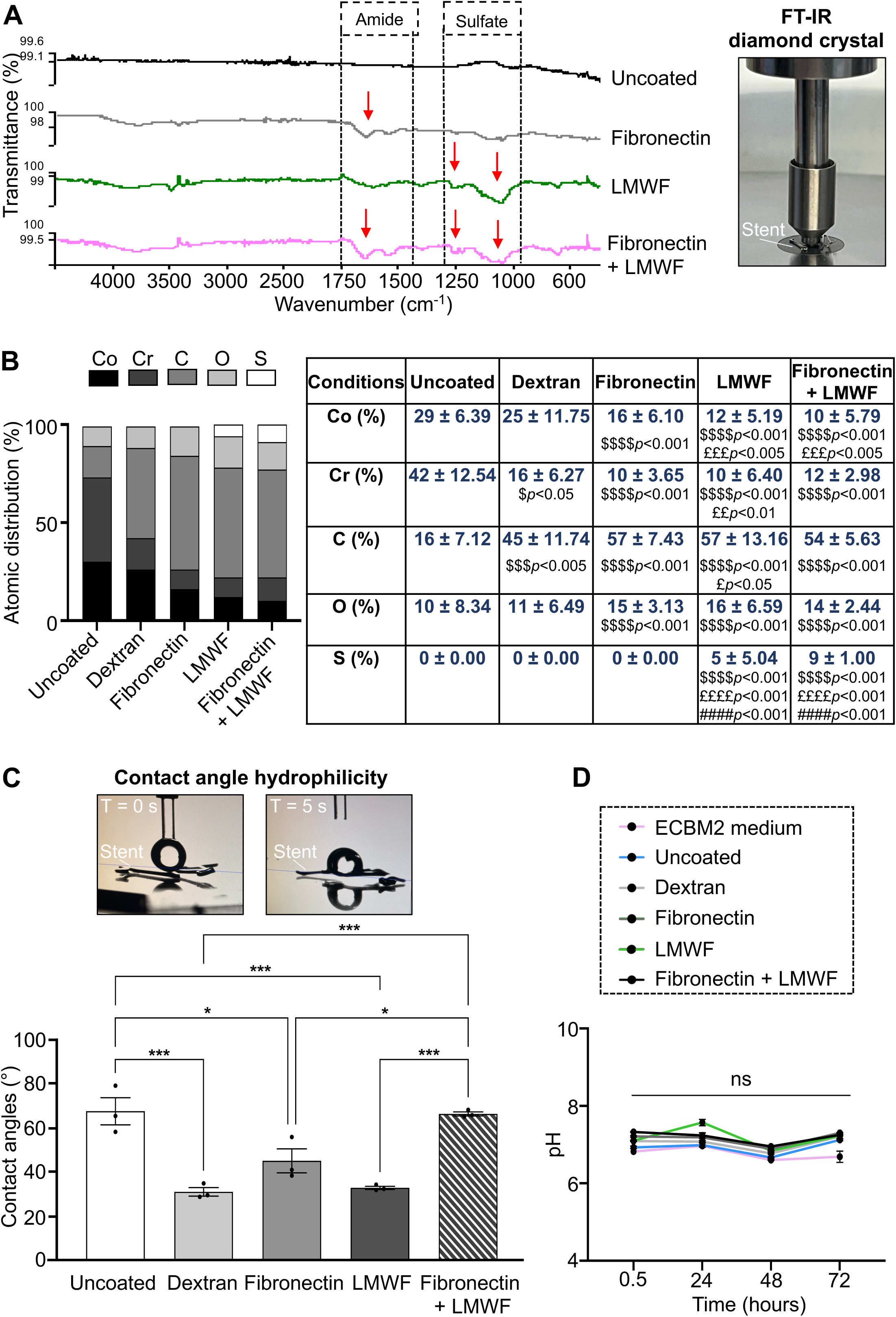
Physicochemical characterization of fibronectin and LMWF coatings on Co-Cr stents. **(A)** Representative Fourier-transform infrared (FT-IR) spectra of uncoated, fibronectin, LMWF and fibronectin-LMWF-coated stents (n ≥ 3). Characteristic absorption bands associated with fibronectin amide groups and LMWF sulfate groups are indicated by red arrows and dashed boxes. **(B)** Energy-dispersive X-ray spectroscopy (EDS) analysis of the surface elemental composition of uncoated, dextran-coated, fibronectin-coated, LMWF-coated and fibronectin-LMWF-coated Co-Cr stents. The stacked bar graph (left panel) shows the relative atomic distribution (%) of cobalt (Co), chromium (Cr), carbon (C), oxygen (O) and sulfur (S) while the table (right panel) reports the quantitative elemental composition (mean ± SEM) obtained from 23 - 60 measurements per condition. Kruskal-Wallis with Dunn’s multiple-comparisons test. Symbols indicate significant differences between conditions: ^$^ *vs* uncoated; ^£^ *vs* dextran; ^#^ *vs* fibronectin; * *vs* LMWF coatings. **(C)** Static water contact angle measurements of uncoated, dextran-coated, fibronectin-coated, LMWF-coated and fibronectin-LMWF-coated Co-Cr stents. Data are presented as mean ± SEM (n = 3). One-way ANOVA with Tukey’s multiple-comparisons test. \**p* < 0.05, \*\**p* < 0.01, \*\*\**p* < 0.005. **(D)** Culture medium pH measured over 72 h in the absence of cells for ECBM2 medium alone or in the presence of uncoated, dextran-coated, fibronectin-coated, LMWF-coated and fibronectin-LMWF-coated stents. Data are mean ± SEM (n = 3). Two-way ANOVA with Tukey’s multiple-comparisons test. ns, not significant.

**Figure 3.**
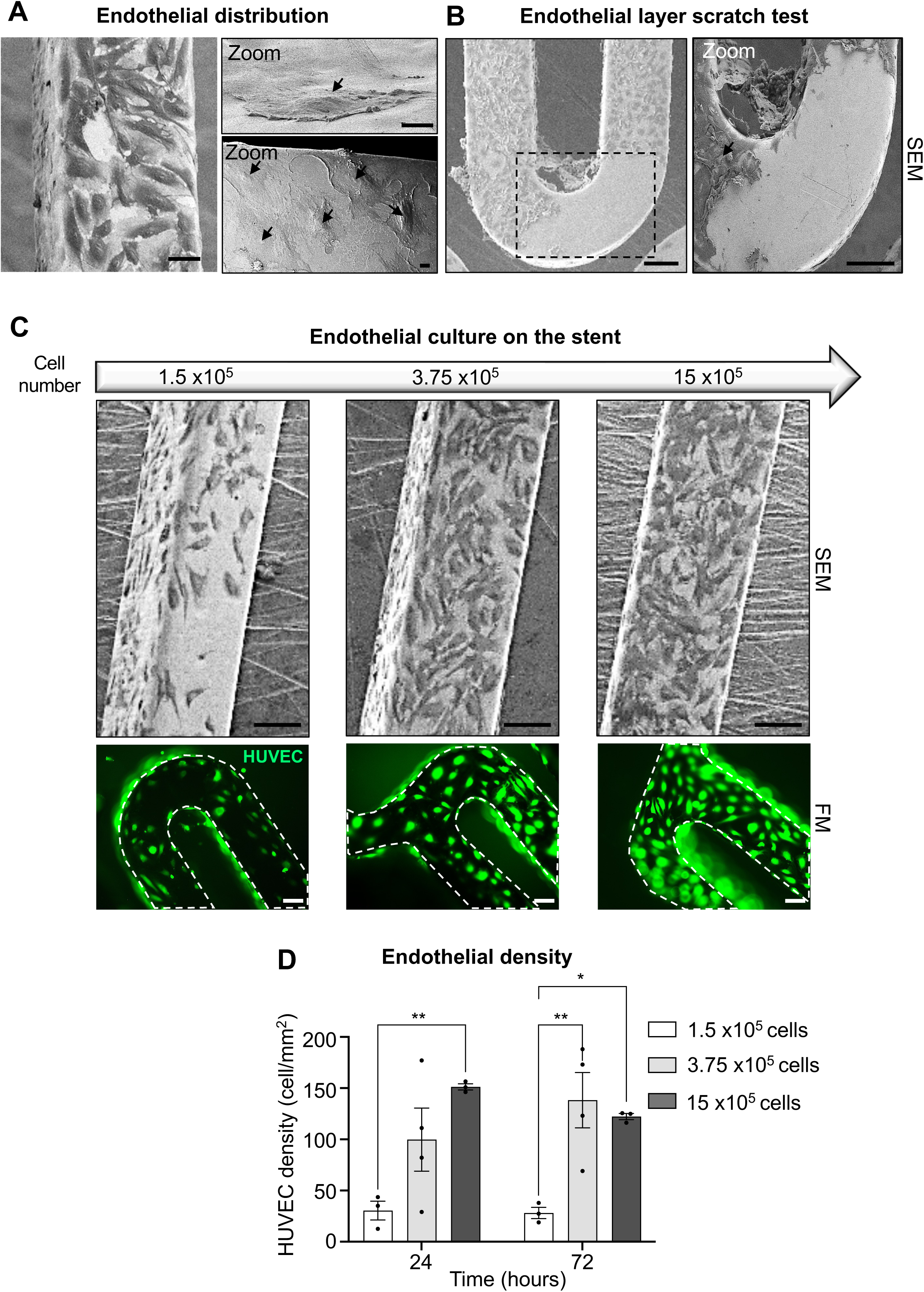
Establishment of an endothelial cell culture model on Co-Cr stents. **(A)** Representative scanning electron microscopy (SEM) images showing the distribution of HUVECs on the upper and lateral surfaces of uncoated Co-Cr stent. The higher-magnification images show HUVECs attached to both surfaces of the stent strut. Scale bars: 100 µm (overview), and 10 µm (higher-magnification, zoom). **(B)** Representative SEM images of an endothelialized uncoated stent following mechanical scratching of the cell layer. The higher-magnification image shows the interface between the remaining endothelial layer and the exposed Co-Cr substrate. Black arrows indicate HUVECs in the higher-magnification images of panels A and B. Scale bars: 100 µm. **(C)** Representative SEM (upper panel) and fluorescence microscopy (FM; lower panel) images of HUVECs cultured for 24 h on uncoated Co-Cr stents using increasing initial cell numbers (1.5 x10^5^, 3.75 x10^5^, and 15 x10^5^ cells). Fluorescence images show CellTracker-labelled HUVECs distributed along the stent struts. White dashed lines delineate the stent boundaries. Scale bars: SEM = 100 µm; FM = 100 µm. **(D)** Quantification of HUVEC density on uncoated stents at 24 and 72 h. Cell density was expressed as the number of HUVECs per mm^2^ of stent surface. Data are mean ± SEM (n ≥ 3 independent experiments). Two-way ANOVA with Tukey’s multiple-comparisons test. The statistical significance is indicated as \**p* < 0.05 and \*\**p* < 0.01.

Fourier-transform infrared spectroscopy (FT-IR) spectra showed distinct features according to the coating composition (Fig. 2A). Fibronectin-coated stents exhibited absorption bands in the amide I and II region (1750 - 1400 cm^-1^). LMWF-coated stents showed absorption in the 1250 - 1000 cm^-1^ region, attributed to S=O (≈ 1250 cm^-1^) and C-O-S (≈ 1050 cm^-1^) stretching vibrations of the sulfate groups consistent withv contributions from the sulfated polysaccharides. Stents coated sequentially with fibronectin and LMWF displayed both spectral features, supporting their presence in the coating. These features were not apparent in the uncoated controls.

Energy-dispersive X-ray spectroscopy (EDS) analysis revealed differences in surface elemental composition between uncoated and coated stents (Fig. 2B). Uncoated stents exhibited cobalt (29.00 ± 6.39 %) and chromium (42.00 ± 12.54 %) as the predominant metallic elements, together with carbon (16.00 ± 7.12 %) and oxygen (10.00 ± 8.34 %); sulfur was not detected. Following dextran coating, cobalt and chromium proportions decreased, while carbon, reflecting the presence of dextran, increased to 45.00 ± 11.74 % (^$$$^*p* < 0.005). No significant differences were observed in oxygen (11.00 ± 6.49 %) compared with uncoated stents.

All coatings reduced the cobalt and chromium signals, with a concomitant increase in carbon, reflecting the deposition of organic material at the stent surface. Dextran coating increased the carbon proportion to 45.00 ± 11.74 % (^$$$^*p* < 0.005), while oxygen (11.00 ± 6.49 %) and sulfur remained unchanged relative to uncoated stents. The protein and polysaccharide coatings had stronger effects: fibronectin, LMWF, and fibronectin-LMWF coatings decreased cobalt to 16.00 ± 6.10 %, 12.00 ± 5.19 %, and 10.00 ± 5.79 %, and chromium to 10.00 ± 3.65 %, 10.00 ± 6.40 %, and 12.00 ± 2.98 %, respectively, while carbon rose to 57.00 ± 7.43 %, 57.00 ± 13.16 %, and 54.00 ± 5.63 %, and oxygen to 15.00 ± 3.13 %, 16.00 ± 6.59 %, and 14.00 ± 2.44 %, respectively (all ^$$$$^*p* < 0.001 *vs* uncoated stents). Sulfur was detected exclusively in LMWF-containing coatings, reaching 5.00 ± 5.04 % after LMWF coating and 9.00 ± 1.00 % after combined fibronectin-LMWF coating (*p* < 0.001 *vs* ^$$$$^ uncoated,^££££^dextran-coated, and ^####^fibronectin-coated stents). Although sulfur content did not differ significantly between these two coatings, its smaller standard deviation with the combined coating (1.00 % vs 5.04 %) SEM suggested a more consistent estimate, potentially reflecting a more homogeneous coating; however, further analysis of inter-stent and intra-stent variability would be required to confirm this hypothesis. Overall, EDS analysis revealed changes in the elemental composition of Cobalt-Chromium (Co-Cr) stent surfaces associated with fibronectin-and fucoidan-containing coatings.

Surface wettability was assessed by static contact angle measurements (Fig. 2C). Uncoated stents exhibited a contact angle of 68.09 ± 7.67°. All individual coatings significantly reduced this value: 31.74 ± 2.55° for dextran (\*\**p* < 0.005), 45.72 ± 7.18° for fibronectin (\**p* < 0.05), and 33.52 ± 0.95° for LMWF (\*\**p* < 0.005), indicating increased surface hydrophilicity. Notably, the hydrophilizing effect of fibronectin was weaker than that of the polysaccharide coatings (\**p* < 0.05 vs LMWF coating). In contrast, the combined fibronectin–LMWF coating did not modify surface wettability: its contact angle (66.91 ± 1.20°) was not significantly different from that of uncoated stents, but was significantly higher than those of all individual coatings (\*\**p* < 0.005 vs dextran and LMWF; \**p* < 0.05 vs fibronectin), indicating that the presence of fibronectin in the coating counteracted the hydrophilizing effect of LMWF. The culture medium pH was monitored over 72 h in the absence of cells (Fig. 2D). No significant differences were observed between coating conditions throughout the incubation period.

Collectively, these results support the successful deposition of fibronectin and LMWF onto Co-Cr stent surfaces. FT-IR spectroscopy and EDS provided complementary evidence of coating deposition, with sulfur detected exclusively in LMWF-containing coatings. Regarding surface properties, while each individual coating significantly increased surface hydrophilicity, fibronectin in the combined coating counteracted the hydrophilizing effect of LMWF, restoring a wettability comparable to that of uncoated stents. Finally, none of the coatings altered the pH of the culture medium, ruling out an effect of coating leaching on subsequent cell-based assays.

### 3.3. Establishment and validation of an endothelial cell culture model on Co-Cr stents

Following coating characterization, an endothelial cell culture model was established on uncoated Co-Cr stents. HUVEC adherence and distribution were first examined by scanning electron microscopy (SEM) (Fig. 3A). Cells covered both the upper and lateral surfaces of the stent struts. Higher-magnification images showed adherent cells extending along the strut surfaces, demonstrating cell attachment across different regions of the stent architecture (Fig. 3A zoom).

To assess the ability of this model to reveal the underlying substrate, a scratch assay was then performed to locally remove the endothelial cell layer and expose the underlying Co-Cr substrate (Fig. 3B). This procedure generated a clearly defined cell-free region adjacent to the remaining endothelial layer, with a distinct interface between the adherent HUVECs and the exposed metallic surface, as shown at higher magnification (Fig. 3B zoom).

At 24h, SEM and fluorescence images showed adherent cells under all three conditions (Fig. 3C). Coverage was sparse at the lowest cell input, and more extensive with 3.75 x10^5^ and 15 x10^5^ cells. However, it did not increase uniformly with the initial cell number.

Cell density was quantified at 24 and 72 h as a function of the initial seeding number (Fig. 3D). At 24 h, density increased with the seeding number, from 30 ± 12 cells/mm² at 1.5 x10^5^ cells to 100 ± 44 cells/mm² at 3.75 x10^5^ cells, reaching 151 ± 4 cells/mm² at 15 x10^5^ cells (\*\**p* < 0.01). At 72 h, cell density reached 138 ± 42 cells/mm² and 122 ± 4 cells/mm² after seeding with 3.75 x10⁵ and 15 x10⁵ cells, respectively, both significantly higher than with 1.5 x10^5^ cells (28 ± 7 cells/mm²; \*\**p* < 0.01 and \**p* < 0.05, respectively). Notably, the highest mean cell density at 72 h was observed with 3.75 x10^5^ cells, suggesting that increasing the seeding number beyond this level did not further improve cell density at this time point.

Based on these results, 3.75 x10^5^ HUVECs per fragment were selected for subsequent experiments, as this seeding number ensured effective early cell attachment and sustained a high cell density at 72 h, while avoiding a higher cell number that provided no further benefit.

### 3.4. Fibronectin-LMWF coating promotes endothelial spreading and lamellipodia formation

Endothelial cell spreading and membrane protrusion formation were subsequently evaluated on uncoated, LMWF-coated and fibronectin-LMWF-coated Co-Cr stents under the previously established seeding conditions. Cell spreading and lamellipodia formation were quantified from fluorescence images acquired at 4 and 72 h, while SEM at 48 h provided complementary morphological assessment of endothelial cell-surface interactions (Fig. 4).

**Figure 4.**
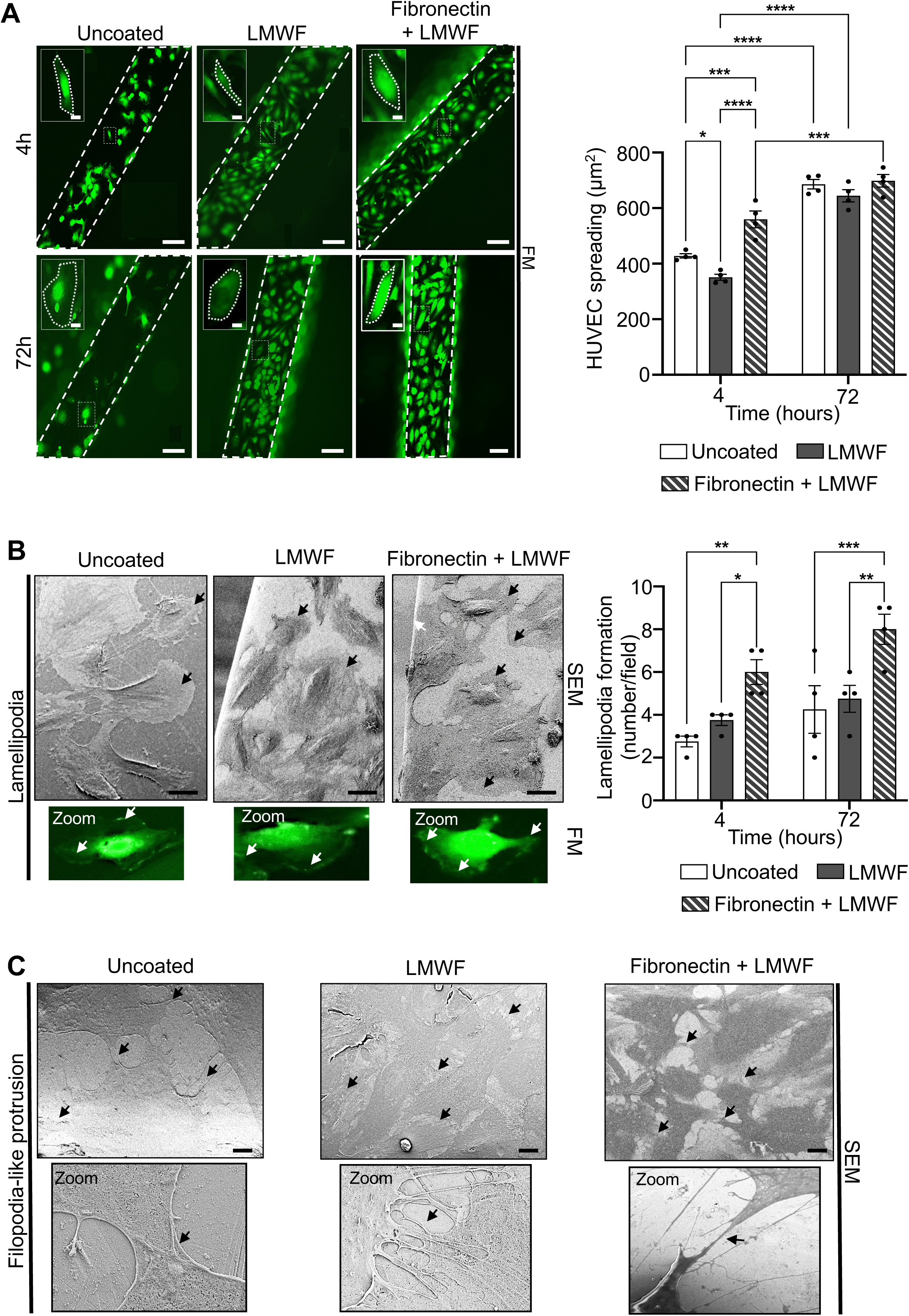
Endothelial spreading and lamellipodia formation on coated Co-Cr stents. **(A)** Representative fluorescence microscopy (FM; left panel) images of HUVECs cultured for 4 and 72 h. Dashed lines mark stent boundaries; boxes indicate higher-magnification insets. HUVECs spreading area (µm²/cell) is quantified on the right. Data are mean ± SEM (n = 4). Two-way ANOVA with Tukey’s multiple-comparisons test. \**p* < 0.05; \*\*\**p* < 0.005; \*\*\*\**p* < 0.001. Scale bars: 100 µm (overview). **(B)** Representative scanning electron microscopy (SEM; left panels) images illustrating lamellipodia formation for 48 h. Higher-magnification FM images show representative lamellipodia (arrows), with quantification per field after 4 and 72 h of culture in the right panels. Data are presented as mean ± SEM (n = 4). Two-way ANOVA with Tukey’s multiple-comparisons test. \**p* < 0.05; \*\**p*< 0.01; \*\*\**p* < 0.005. Scale bars: SEM = 30 µm. **(C)** Representative SEM images illustrating filopodia-like protrusion formation for 48 h. High-magnification images show selected regions, with arrows indicating representative filopodia. Filopodia were qualitatively assessed but not quantified. Scale bars: 20 µm.

Cell spreading was quantified as the projected cell area from fluorescence images (Fig. 4A). At 4 h, HUVECs on fibronectin-LMWF-coated stents exhibited a significantly larger cell area (559.52 ± 45.00 µm²) than those on uncoated (427.99 ± 11.00 µm², \*\*\**p* < 0.005) and LMWF-coated stents (350.98 ± 17.00 µm², \*\*\*\**p* < 0.001), indicating enhanced early cell spreading on the combined coating. From 4 to 72h, cell area increased significantly within each surface condition, reaching 685.94 ± 29.00 µm² on uncoated stents, 644.17 ± 33.00 µm² on LMWF-coated stents, and 698.18 ± 31.00 µm² on fibronectin-LMWF-coated stents (within-group comparisons *vs* 4 h: \*\*\*\**p* < 0.001 and \*\*\**p* < 0.005). As a result, cell area no longer differed significantly among the three surfaces at 72 h.

Furthermore, representative SEM images revealed lamellipodia and filopodia at the cell periphery under all three conditions (Fig. 4B, left panels and 4C). Lamellipodia formation was quantified from the corresponding FM images (insets, Fig. 4B, right panels). At 4 h, fibronectin-LMWF coated stents showed 6 ± 0.87 lamellipodia formation per field, significantly more than the number observed on uncoated stents (3.00 ± 0.47 per field; \*\**p* < 0.01) and LMWF-coated stents (4.00 ± 0.36 per field; \**p* < 0.05). This difference persisted at 72 h: fibronectin-LMWF-coated stents exhibited 8.00 ± 0.94 lamellipodia per field, significantly more than the values observed on uncoated stents (4.00 ± 1.82, \*\*\**p* < 0.005) and LMWF-coated stents (5.00 ± 1.00, \*\**p* < 0.01). At 48 h, SEM images confirmed the presence of filopodia-like protrusions on all three surfaces, and higher-magnification insets highlighted their morphology (Fig. 4C). These structures were assessed qualitatively and not quantified.

Taken together, these findings suggest that fibronectin-LMWF coating enhanced early endothelial cell spreading and increased lamellipodia formation per field at both 4 and 72 h, although differences in cell area were no longer significant at 72 h. Filopodia-like protrusions were observed under all three surface conditions.

### 3.5. Fibronectin-LMWF coating supports endothelial colonization and proliferative activity on Co-Cr stents

Following the analyses of spreading, lamellipodia and filopodia, we examined cell colonization over time and proliferative activity. Lactate dehydrogenase (LDH) release was assessed in parallel to evaluate cell membrane damage under the different coating conditions (Fig. 5).

**Figure 5.**
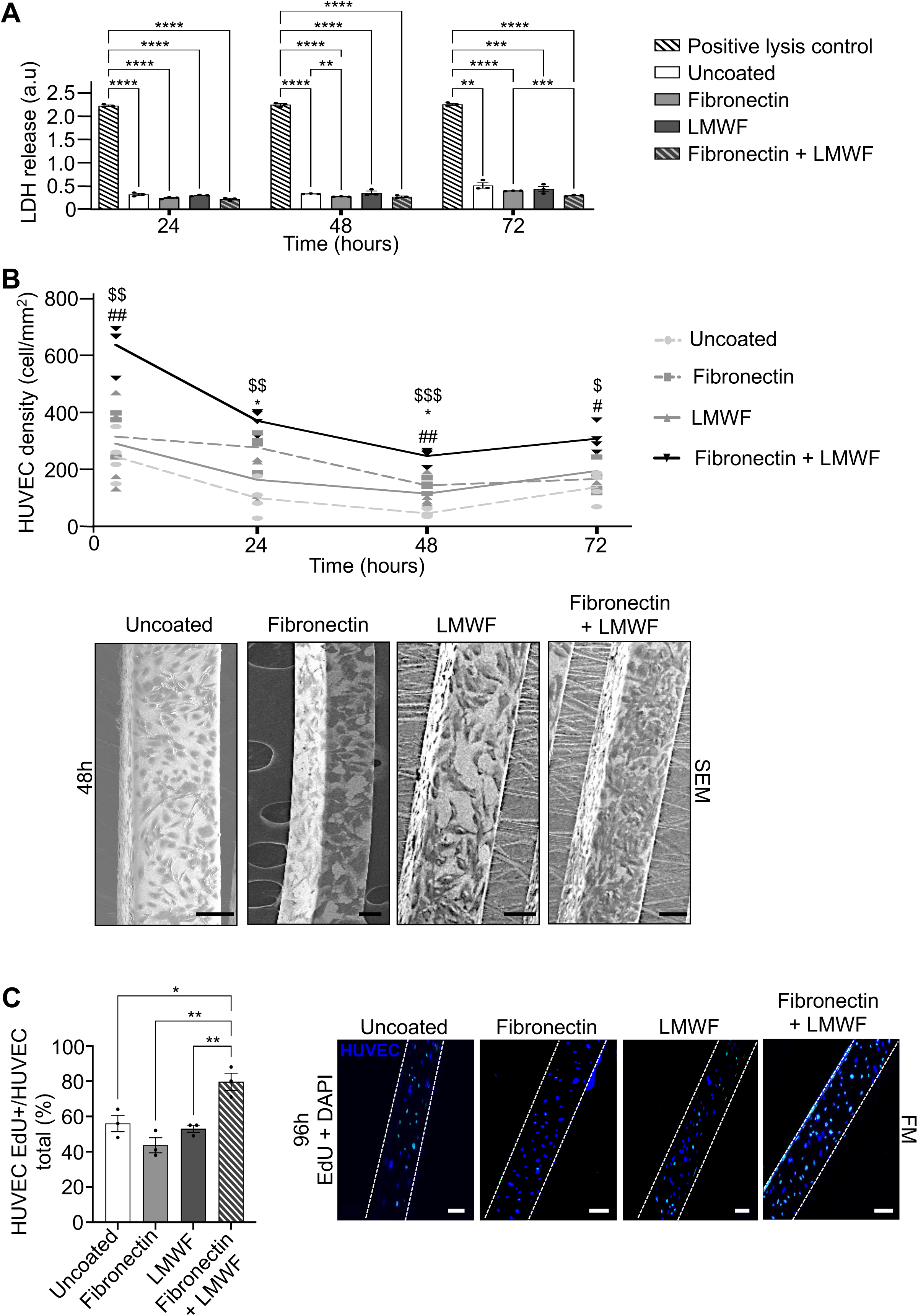
Endothelial cytocompatibility and proliferation on Co-Cr stents with surface coatings. **(A)** Lactate dehydrogenase (LDH) released from HUVEC cultured on uncoated, fibronectin-coated, LMWF-coated and fibronectin-LMWF-coated stents after 24 h, 48 h and 72 h. Positive lysis control (permeabilized HUVECs) was included as an assay control. Data are mean ± SEM (n = 3). Two-way ANOVA with Tukey’s multiple-comparisons test. \*\**p* < 0.01; \*\*\**p* < 0.005; \*\*\*\**p* < 0.001. **(B)** Representative scanning electron microscopy (SEM) images of HUVECs cultured for 48 h on the different stent surfaces, illustrating endothelial cell coverage along the stent struts (lower panel). Scale bars: 100 µm. HUVEC density was quantified at 4, 24, 48 and 72 h after seeding and expressed as cells/mm^2^ (upper panel). Data are mean ± SEM (n = 4). Two-way ANOVA with Tukey’s multiple-comparisons test. Symbols indicate significant differences for fibronectin-LMWF-coated stents *vs* uncoated ($), fibronectin-coated (#) or LMWF-coated (*) stents (^$,#,*^*p* < 0.05; ^$$^,^##^*p* < 0.01; ^$$$^*p* < 0.005). **(C)** Representative merged fluorescence microscopy (FM) images of HUVECs at 96 h, showing EdU staining (green) and DAPI nuclear counterstaining (blue) (right). White dashed lines delineate the stent boundaries. Scale bars: 100 µm. HUVEC proliferation was assessed by EdU incorporation between 72 h and 96 h after seeding and quantified as the percentage of EdU-positive HUVECs relative to the total number of HUVEC population (left). Data are mean ± SEM (n = 3). One-way ANOVA with Tukey’s multiple-comparisons test. \**p* < 0.05; \*\**p* < 0.01.

The cytocompatibility of the different biomolecular coatings was assessed by measuring LDH release after 24, 48, and 72 h of culture, relative to the positive lysis control (permeabilized cells; 2.23 ± 0.02, 2.25 ± 0.03, and 2.36 ± 0.03 at 24, 48, and 72 h, respectively; Fig. 5A). At all three time points, LDH levels remained far below the positive control for every surface condition, indicating limited cell damage (\*\*\*\**p* < 0.001 for all comparisons at 24 and 48 h; \*\**p* < 0.01 to \*\*\*\**p* < 0.001 at 72 h). On uncoated stents, LDH levels were 0.32 ± 0.03, 0.33 ± 0.00, and 0.51 ± 0.08 at 24, 48, and 72 h, respectively. Fibronectin-coated stents showed 0.24 ± 0.02, 0.27 ± 0.00, and 0.40 ± 0.00; LMWF-coated stents 0.30 ± 0.01, 0.35 ± 0.05, and 0.44 ± 0.07; and fibronectin–LMWF-coated stents 0.22 ± 0.02, 0.27 ± 0.02, and 0.30 ± 0.01.

Beyond this global cytocompatibility, coating-dependent differences emerged. At 24 h and 48 h, fibronectin-LMWF-coated stents exhibited lower LDH release than uncoated stents (**** *p* < 0.001, Fig. 5A). At 48 h, fibronectin-coated stents also showed lower LDH release than uncoated stents (\*\**p* < 0.01). At 72 h, fibronectin-LMWF-coated stents showed significantly lower LDH levels than fibronectin-coated stents (\*\*\**p* < 0.005).

Taken together, these results indicate that all coatings were cytocompatible over the 72 h culture period, and that the fibronectin-LMWF coating limited LDH release at every time point significantly more than uncoated stents at 24 and 48 h, and more than fibronectin-coated stents at 72 h.

Endothelial cell density was quantified at 4, 24, 48, and 72 h after cell seeding (Fig. 5B, lower panel). Fibronectin-LMWF-coated stents supported the highest cell density at every time point.

At 4 h, cell density on fibronectin-LMWF-coated stents reached 638.00 ± 58.50 cells/mm², significantly higher than on uncoated (245.00 ± 61.00; ^$$^*p* < 0.01) and fibronectin-coated stents (315.00 ± 71.50; ^##^*p* < 0.01), but comparable to LMWF-coated stents. At 24 h, cell density decreased in all groups, yet fibronectin-LMWF-coated stents maintained a density of 371.50 ± 28.50 cells/mm², significantly higher than on uncoated (99.75 ± 44.25; ^$$^*p* < 0.01) and LMWF-coated stents (164.25 ± 66.75; \**p* < 0.05). At 48 h, this advantage extended to all three comparators: 247.50 ± 20.70 cells/mm² on fibronectin-LMWF-coated stents, versus 45.75 ± 8.62 on uncoated (^$$$^*p* < 0.001), 144.00 ± 16.00 on fibronectin-coated (^##^*p* < 0.01), and 115.25 ± 38.37 on LMWF-coated stents (\**p* < 0.05). At 72 h, fibronectin-LMWF-coated stents still exhibited the highest density (308.25 ± 32.87) significantly higher than uncoated (138.25 ± 42.25; ^$^*p* < 0.05) and fibronectin-coated stents (167.25 ± 41.75; ^#^*p* < 0.05), but comparable to LMWF-coated stents (194.75 ± 53.25).

Notably, the density profile of fibronectin-LMWF-coated stents followed a distinct pattern over time, with an initial peak at 4 h (638.00 ± 58.50 cells/mm²) followed by a progressive decrease until 48 h (247.50 ± 20.70 cells/mm²) and a slight increase at 72 h (308.25 ± 32.87 cells/mm²). Importantly, cell density remained significantly higher on fibronectin-LMWF-coated stents than on fibronectin-coated stents at all time points, indicating that the addition of LMWF to fibronectin was associated with significantly greater endothelial cell density throughout the study period.

Scanning electron microscopy (SEM) images acquired at 48 h showed adherent HUVECs along the stent struts under all four conditions (Fig. 5B, lower panels). Cells displayed spread morphologies, with differences in the extent and distribution of surface coverage. Complementary fluorescence images illustrated HUVEC distribution at 4, 48 and 72 h (Supplementary Fig. 5).

Finally, HUVEC proliferation was assessed by 5-ethynyl-2′-deoxyuridine (EdU) incorporation from 72 h to 96 h after seeding. Representative FM images at 96 h showed EdU-positive nuclei under all four conditions (Fig. 5C, right panel). Quantification revealed a significantly higher proliferation rate on fibronectin-LMWF-coated stents (79.67 ± 5.78 %) than on uncoated (56.00 ± 5.33 %, \**p* < 0.05), fibronectin-coated (43.67 ± 5.56 %, \*\**p* < 0.01) and LMWF-coated stents (53.00 ± 2.67 %, \*\**p* < 0.01) (Fig. 5C, left panel).

Collectively, these findings indicate that the fibronectin-LMWF coating stands out on all three counts: it supported the highest mean cell density throughout the observation period, elicited the strongest proliferative response between 72 and 96 h. Regardless of the coating type, LDH release remained low, indicating limited membrane damage and no evidence of substantial cytotoxicity.

### 3.6. Junctional, cytoskeletal, and adhesion-associated protein organization in endothelial cells on coated Co-Cr stents

We next investigated how the different coatings influence the endothelial phenotype after 48 h of culture, by assessing endothelial junctional organization (cell-cell), cytoskeletal architecture and cell-matrix adhesion-associated proteins. Immunostaining for VE-cadherin (junctions), F-actin (cytoskeleton), integrin β1 and α6, and syndecans-1 and-4 (SDC-1 and SDC-4; adhesion-associated proteins) provided spatial information on their cellular localization (Fig. 6), while Western blotting assess the abundance of VE-cadherin and integrins, as well as the phospho-paxillin/total paxillin ratio (Supplementary Fig. 6). Complementary morphological analysis was performed by SEM (Fig. 6B lower panel).

**Figure 6.**
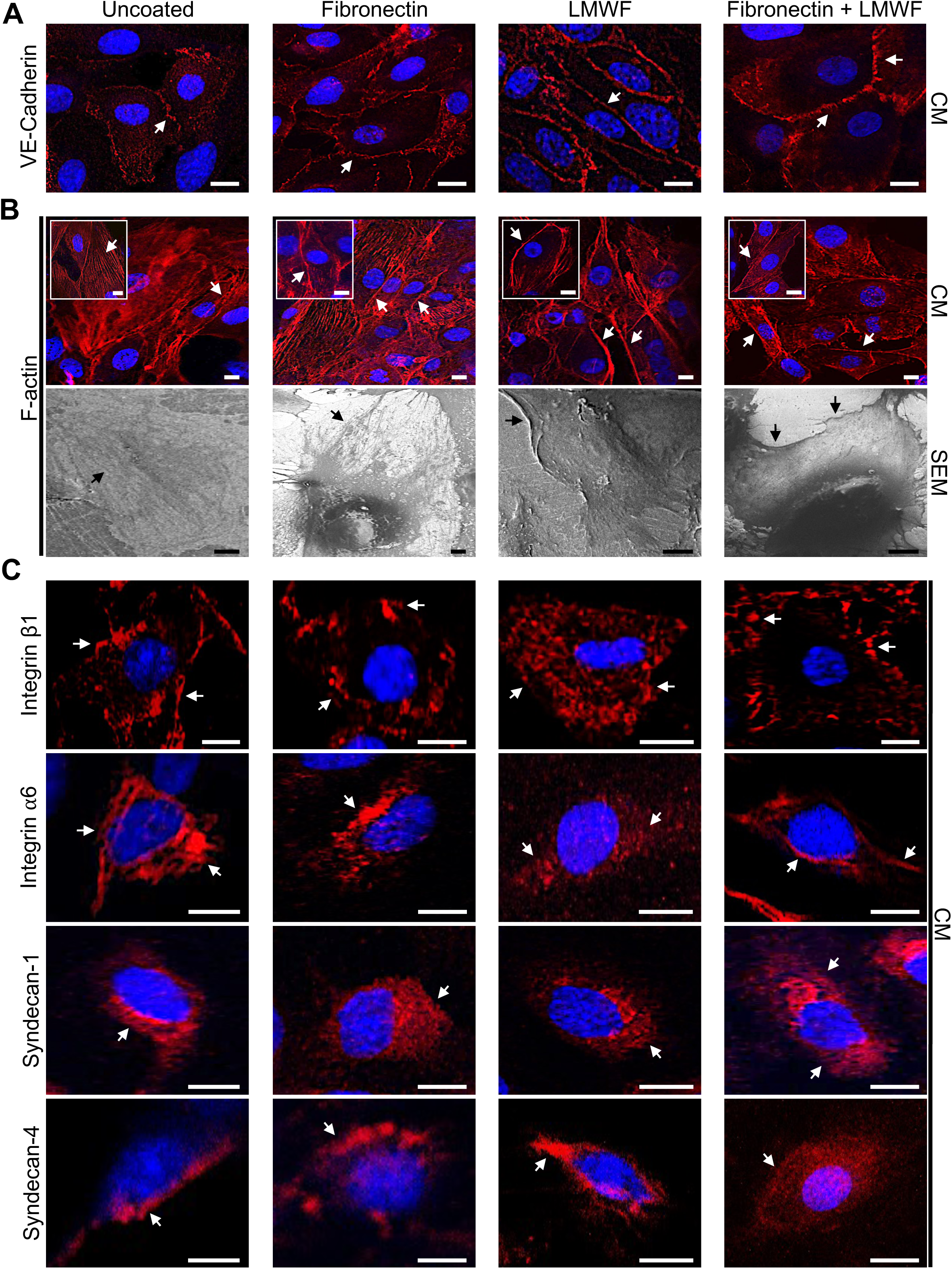
Organization of endothelial junctional, cytoskeletal, and adhesion-associated proteins on coated Co-Cr stents. Representative images of HUVECs cultured for 48 h on uncoated, fibronectin-coated, LMWF-coated and fibronectin-LMWF-coated Co-Cr stents. In all confocal microscopy (CM) images, the indicated target protein is shown in red, and nuclei are counterstained with DAPI (blue). **(A)** CM images showing VE-Cadherin distribution. Arrows indicate representative staining at cell-cell junctions. Scale bars: 30 µm. **(B)** CM (upper panels) and scanning electron microscopy (SEM; lower panels) images showing F-actin organization and cell morphology, respectively. F-actin was visualized using Alexa Fluor 568-conjugated phalloidin. Arrows indicate representative submembranous and cytoplasmic F-actin staining patterns. Inserts show higher-magnification views of representative cells, illustrating the corresponding F-actin distribution patterns. Scale bars: CM = 30 µm and SEM = 10 µm. **(C)** CM images showing integrin β1, integrin α6, syndecan-1 (SDC-1) and syndecan-4 (SDC-4) distribution. Arrows indicate representative membrane and cytoplasmic localization. Scale bars: 30 µm.

VE-cadherin staining was detected at cell-cell contacts under all four surface conditions (Fig. 6A). Representative images suggested distinct junctional staining patterns across conditions: LMWF-coated stents exhibited a more regular distribution, whereas fibronectin-LMWF-coated stents displayed thicker and more intense staining along some cell-cell interfaces. Thus, the coatings appeared to influence the spatial organization of VE-cadherin rather than its total abundance, as confirmed by Western blotting analysis revealing no significant differences between conditions (Supplementary Fig. 6A).

F-actin staining revealed differences in cytoskeletal organization across the representative images (Fig. 6B, upper panel). On uncoated and fibronectin-coated stents, HUVECs displayed prominent filamentous bundles distributed throughout the cell body, with fibers predominantly aligned in parallel. In contrast, on LMWF-coated and fibronectin-LMWF-coated stents, F-actin appeared more concentrated in the submembranous region, as particularly evident in the higher-magnification images. These distinct observations suggest that coating may influence cytoskeletal organization in association with cell-cell junctional organization, although whether these changes reflect enhanced intercellular interactions remains to be established through further quantitative analysis. SEM images provided complementary information on cell morphology and membrane extensions (Fig. 6B, lower panel).

The distribution of cell-matrix adhesion-associated proteins was examined by confocal microscopy (Fig. 6C, first row). Integrin β1 staining was detected throughout the cell body and along the cell periphery, with distinct patterns across surface conditions: predominantly punctate, with both peripheral and intracellular labeling, on uncoated and fibronectin-coated stents; more homogeneous and cytoplasmic on LMWF-coated stents; and more pronounced at the plasma membrane on fibronectin-LMWF-coated stents. However, Western blotting analysis detected no significant differences in total integrin β1 expression between conditions (Supplementary Fig. 6B) suggesting that coatings influenced the spatial organization of integrin β1 rather than its total abundance.

Integrin α6 also displayed distinct staining patterns across conditions (Fig. 6C, second row). On uncoated and fibronectin-coated stents, staining was predominantly perinuclear and cytoplasmic, extending toward the cell periphery. On LMWF-coated stents, the signal appeared more diffuse, throughout the cytoplasm, whereas fibronectin-LMWF-coated stents exhibited more pronounced peripheral and linear staining. At the protein level, total integrin α6 abundance did not differ significantly between conditions (Supplementary Fig. 6C).

SDC-1 staining was mainly perinuclear on uncoated, fibronectin-coated, and LMWF-coated stents (Fig. 6C, third row). In contrast, HUVECs cultured on fibronectin-LMWF-coated stents showed broader cytoplasmic staining extending beyond the perinuclear region. Distinct staining patterns were also observed for SDC-4 (Fig. 6C, fourth row): peripheral labeling on uncoated stents, both peripheral and punctate intracellular staining on Fibronectin-coated stents; more pronounced, with punctate staining along the cell border on LMWF coated stents; and broader cytoplasmic staining on fibronectin-LMWF coated stents. Notably, the two syndecans showed a shared pattern: perinuclear or peripheral localization on all surfaces except the fibronectin-LMWF coating, where staining extended throughout the cytoplasm. The phospho-paxillin (Tyr118)/total paxillin ratio showed no significant differences between conditions (Supplementary Fig. 6D).

Overall, surface modification altered the spatial organization of endothelial junctional, cytoskeletal, and cell-matrix adhesion-associated proteins. Differences in VE-cadherin and F-actin localization, together with the redistribution of integrin β1, integrin α6, SDC-1, and SDC-4, were observed across the four surface conditions. Importantly, these redistributions occurred without significant differences in total protein abundance (VE-cadherin, integrin β1, α6; supplementary Fig. 6A-C), suggesting that surface coatings may modulate protein distribution rather than overall expression. Further quantitative analyses are needed to determine the functional implications of these spatial changes for endothelial organization and adhesion.

### 3.7. Endothelial soluble mediators and smooth muscle cell responses to conditioned media

Following the assessment of endothelial colonization and proliferation, we next examined the paracrine activity of the endothelial layer. We profiled soluble mediators released by HUVECs using a multiplex immunoassay and investigated the effects of HUVEC-conditioned media on smooth muscle cell (SMC) migration and metabolic activity (Fig. 7). For multiplex analysis, HUVECs were first cultured in complete medium for 24 h, after which the medium was replaced with serum-free medium. Conditioned media were collected after a further 24 or 48 h of cell culture.

**Figure 7.**
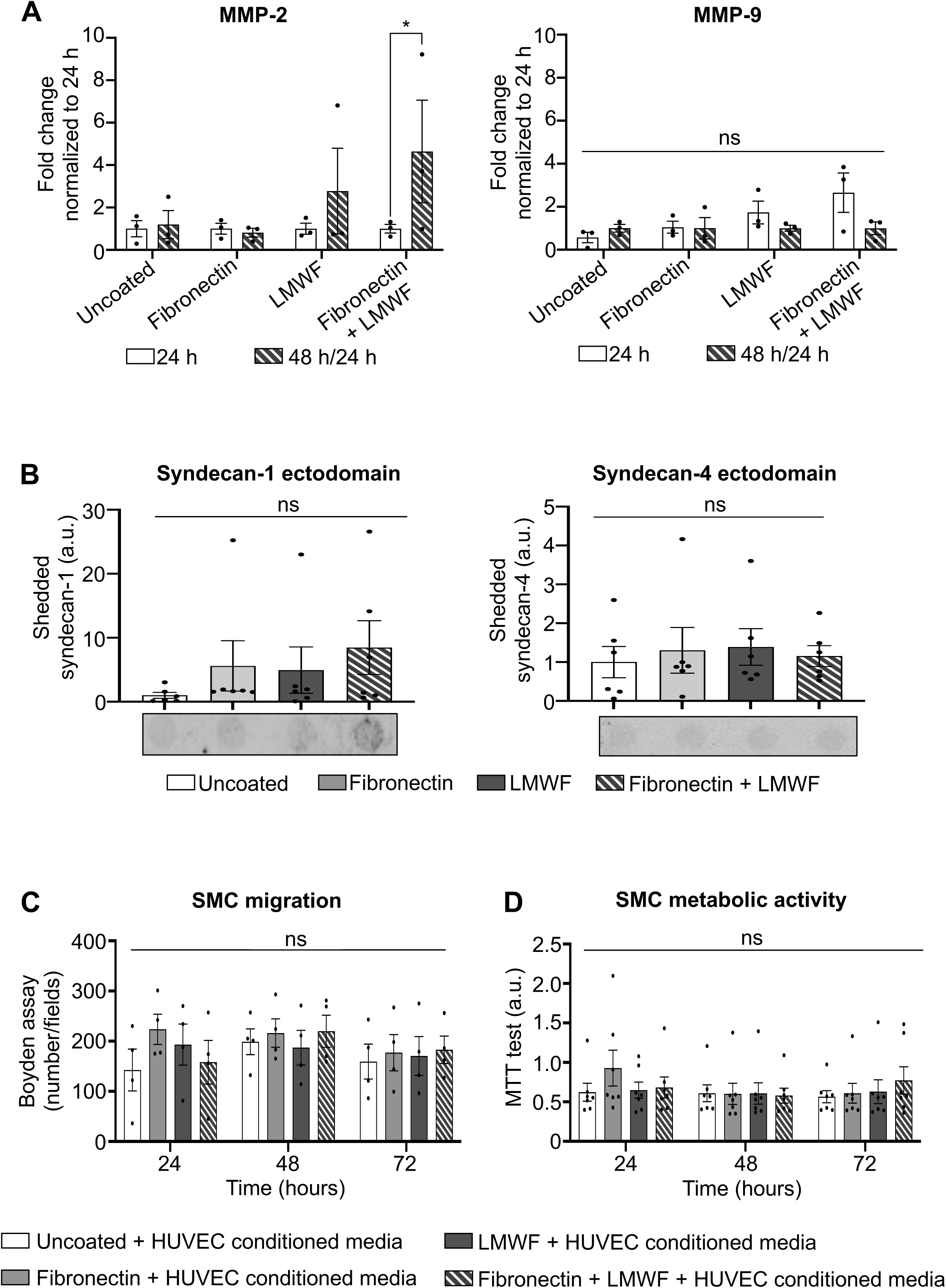
Release of MMP and SDC ectodomains in conditioned HUVEC media and its effects on SMC migration and viability. **(A)** Quantification of MMP-2 (left) and MMP-9 (right) fold changes in culture supernatants from HUVEC cultured on uncoated, fibronectin-coated, LMWF-coated, and fibronectin-LMWF-coated Co-Cr stents. Cells were cultured for 24 h in complete medium, followed by 24 and 48 h in serum-free medium before supernatant collection. Data are presented as mean ± SEM (n = 3). Kruskal-Wallis test for MMP-2 and one way ANOVA for MMP-9, followed by Dunn’s and Tukey’s multiple-comparisons test, respectively. \**p* < 0.05; ns, not significant. **(B)** Representative Dot blot and quantification of soluble syndecan-1 (SDC-1, left) and syndecan-4 (SDC-4, right) ectodomains in conditioned media collected after 48h. Signal intensities were normalized to the uncoated condition (arbitrary units, a.u.). Data are mean ± SEM (n = 6). One-way ANOVA with Tukey’s multiple-comparisons test. **(C)** SMC migration in response to conditioned media collected from HUVECs cultured on uncoated, fibronectin-coated, LMWF-coated, and fibronectin-LMWF-coated stents, assessed using a Boyden chamber assay at 24, 48 and 72 h. Migration is expressed as the number of migrated SMCs per field. Data are mean ± SEM (n = 4). Two-way ANOVA with Tukey’s multiple-comparisons test. **(D)** SMC metabolic activity following exposure to HUVEC-conditioned media, assessed by MTT assay at 24, 48, and 72 h. Data are mean ± SEM (n = 3). Two-way ANOVA with Tukey’s multiple-comparisons test.

Matrix metalloproteinase-2 and-9 (MMP-2 and MMP-9) concentrations were quantified as indicators of matrix-remodeling activity (Fig. 7A). Temporal changes were expressed as fold changes, calculated as the ratio of MMP concentrations at 48 h to those at 24 h for each biological replicate. Notably, MMP-2 increased significantly in the fibronectin-LMWF condition (4.63 ± 3.05; \**p* < 0.05), whereas no significant temporal changes were observed in the other coating conditions. MMP-9 fold changes did not differ significantly across surface conditions.

Soluble SDC-1 and SDC-4 ectodomains released by shedding were quantified by dot blot analysis (Fig. 7B). Normalized signal intensities did not differ significantly across surface conditions, although substantial inter-sample variability was observed, particularly in the coated groups.

As our stent surface coating influenced endothelialization, and endothelial cells can in turn modulate SMC proliferation, we investigated whether the endothelial response to the different coatings affected SMC behavior. This is particularly relevant *in vivo*, where limiting excessive SMC proliferation is important to prevent intimal hyperplasia after stent implantation. The effects of HUVEC-conditioned media on SMC behavior were assessed at 24, 48 and 72 h. Neither Boyden chamber assays (Fig. 7C) nor MTT metabolic activity (Fig. 7D) revealed significant differences between the four surface conditions at any time point.

Inflammation and angiogenesis are closely linked to endothelialization and vascular healing. Therefore, we quantified growth factors involved in angiogenic responses (VEGF-A and FGF-2), chemokines associated with inflammatory signaling and cell recruitment (MCP-1/CCL2, RANTES/CCL5 and IL-8/CXCL8), and cytokines involved in the regulation of inflammation (IL-6, TNF-α and IL-10) to further characterize the endothelial secretory profile (Supplementary Fig. 7). Significant temporal increases were observed in the uncoated condition for both VEGF-A, which rose from 93.40 ± 11.02 pg/mL at 24 h to 130.66 ± 26.63 pg/mL at 48 h (\*\**p* < 0.01; Supplementary Fig. 7A), and IL-6, which increased from 2.68 x10^2^ ± 0.27 x10^2^ to 4.20 x10^2^ ± 1.48 x10^2^ pg/mL over the same period of serum-free culture (\**p* < 0.05; Supplementary Fig. 7F). No significant differences in TNF-α, IL-10, FGF-2, MCP-1/CCL2, or RANTES/CCL5 concentrations were observed across surface conditions (Supplementary Fig. 7B–D, G–H).

Interestingly, IL-8/CXCL8 concentrations increased significantly between 24 and 48 h of serum-free culture in the uncoated, LMWF-coated, and fibronectin-LMWF-coated conditions (all \**p* < 0.05; Supplementary Fig. 7E). Concentrations increased from 1.26 x10^4^ ± 0.33 x10^4^ to 2.08 x10^4^ ± 0.85 x10^4^ pg/mL in the uncoated condition, from 0.81 x10^4^ ± 0.08 x10^4^ to 1.64 x10^4^ ± 0.49 x10^4^ pg/mL in the LMWF-coated, and from 0.87 x10^4^ ± 0.17 x10^4^ to 1.78 x10^4^ ± 0.23 x10^4^ pg/mL in the fibronectin-LMWF-coated condition.

Together, these results revealed an increased 48 h/24 h fold change in MMP-2 concentration in the fibronectin-LMWF condition, while soluble syndecan signals and the migration and metabolic activity of SMCs exposed to HUVEC-conditioned media did not differ significantly between surface conditions. Additional mediator analyses identified temporal changes in IL-8/CXCL8 concentrations in the LMWF-coated and fibronectin-LMWF-coated conditions, but did not establish a distinct inflammatory profile associated with the combined coating.

## 4. Discussion and perspectives

### ❖ Fibronectin-assisted LMWF coating creates an interface that supports endothelial colonization

This study advances cardiovascular biomaterials research by systematically evaluating the physicochemical properties and endothelial responses associated with fibronectin-LMWF coating directly on Co-Cr stent struts, a challenging geometry for surface modification and biological assessment. Fibronectin precoating enhanced LMWF deposition and retention, with an approximately ten-fold higher initial surface-associated LMWF-DTAF fluorescence compared with LMWF-DTAF coating alone, maintained throughout the 7 days study. Importantly, this modification was accompanied by improved early cell spreading. Throughout the observed period, this surface treatment improved lamellipodia formation, higher endothelial cell density and increased proliferation. These findings establish the feasibility of fibronectin-LMWF coating directly on Co-Cr stent struts and support its potential to promote endothelial colonization *in vitro*. The clinical use of Optimus-L Co-Cr stents in infants and young children with congenital heart disease provides a relevant application for investigating surface-modification strategies. In this population, maintaining vascular patency is particularly important, and improved endothelial cell colonization may represent one avenue for optimizing the biological performance of metallic stents. Further studies are needed to establish whether these *in vitro* effects translate into improved vascular healing and long-term patency [23,24,25]. These potential benefits may also extend to other Co-Cr stent platforms used to treat coronary and small peripheral artery disease, supporting the broader applicability of surface modification strategies across vascular interventions.

Complementary surface-characterization methods provided converging evidence of successful coating deposition. Fluorescence microscopy showed the distribution of LMWF-DTAF across the stent struts, while FT-IR provided information on identifying the chemical groups present in the surface-associated material. The interpretation of protein-associated infrared bands can be informed by previous work characterizing the amide I and II regions [30]. In addition, energy-dispersive X-ray spectroscopy (EDS) detected sulfur on LMWF-containing surfaces, while no sulfur was detected on the non-sulfated dextran control, confirming that the EDS signal was attributable to the sulfated polysaccharide. Considered alongside the fluorescence-retention measurements, these findings support successful surface modification. Nevertheless, these techniques do not independently establish the molecular interactions responsible for enhanced LMWF retention, and their interpretation should take into account the relevant control surfaces and the sensitivity and specificity of each method.

The enhanced LMWF retention following fibronectin precoating may reflect interactions between LMWF and the adsorbed protein layer. Fibronectin adsorption is influenced by surface chemistry, which can affect protein adsorption and its subsequent biological presentation [31,32]. We hypothesize that fibronectin adsorbs onto the Co-Cr surface through hydrophobic interactions, potentially exposing positively charged regions, including lysine-and arginine-rich sequences, that may interact electrostatically with the negatively charged sulfate groups of LMWF. Fibronectin may thus act as a scaffold linker between the metallic Co-Cr stent and the polysaccharide LMWF. Previous studies provide a rationale for the proposed interaction between fucoidan and fibronectin. Lyon *et al*. showed that heparan sulfate binding to the Hep-II domain of fibronectin depends on structural features, including sulfation patterns and oligosaccharide length [20]. More directly, Liu *et al*. reported that fucoidan binds to fibronectin at multiple sites, including heparin-binding sequences, and that this interaction is sensitive to pH and ionic strength-dependent interactions involving multiple sites along the protein [12]. Although interactions between sulfated polysaccharides and protein-binding domains are well documented [33,34], the molecular interactions underlying enhanced LMWF retention remain to be elucidated.

In the present study, enhanced LMWF retention following fibronectin precoating was associated with favorable endothelial responses at selected time points, suggesting that protein-polysaccharide interactions may contribute to improved endothelialization of Co-Cr stent surfaces. Supporting this hypothesis, we previously demonstrated that fucoidans bind to laminin through dose-dependent, saturable interactions involving binding sites distinct from those involved in the binding of heparin and other glycosaminoglycans (GAGs), including chondroitin sulfate, as well as dextran sulfate, a sulfated dextran derivative [35]. We also demonstrated the affinity of radiolabeled fucoidan for P-selectin expressed by activated platelets and endothelial cells, supporting its potential use for imaging endothelial activation [36]. Collectively, these findings highlight the capacity of fucoidan to engage in protein-specific interactions, whose biological consequences depend on the protein and experimental context. By analogy, specific interactions between LMWF and the precoated fibronectin layer may contribute to both the enhanced LMWF retention and the favorable endothelial responses observed here, although the molecular basis of these interactions remains to be clarified.

Fucoidan-protein interactions have also been investigated in endothelialization of vascular biomaterials. We previously demonstrated that fucoidan-VEGF-based polyelectrolyte multilayer coating improved endothelial cell adhesion and colonization of decellularized porcine pulmonary valves [17]. Similarly, Yao *et al.* reported that fucoidan functionalization combined with topographical modification of acellular synthetic vascular grafts promoted fibronectin recruitment and supported integrin-associated adhesion and focal-adhesion formation [37]. Although these coating architectures differ from the sequential fibronectin-LMWF application directly to Co-Cr stent struts in the present study, they support the relevance of fucoidan-based surface engineering and its interplay with adhesive proteins. Whether integrin-mediated mechanisms contribute to the endothelial responses observed here remains to be established.

Importantly, fucoidan-protein interactions do not necessarily promote cell adhesion in all biological contexts. Liu *et al.* reported that fucoidan inhibited adenocarcinoma cell adhesion to fibronectin [12], illustrating that the biological outcome depends on cell type, protein presentation, and material environment. Thus, the endothelial responses observed here should be interpreted in the context of the specific fibronectin-LMWF coating architecture rather than as a universal effect of fucoidan. Further studies are needed to elucidate the molecular interactions responsible for enhanced LMWF retention and their contribution to endothelial adhesion and colonization on Co-Cr stent surfaces.

Notably, the fibronectin-LMWF coating elicited the most favorable endothelial responses, including enhanced cell spreading, lamellipodia formation, as confirmed by quantitative analysis, and proliferation, while largely preserving the wettability of bare Co-Cr (water contact angles: 66.91 ± 1.20° and 68.09 ± 7.67°, respectively). In contrast, LMWF-coated, and fibronectin-coated surfaces exhibited increased wettability. Interestingly, the combined fibronectin-LMWF coating exhibited a significantly higher water contact angle than the individual coatings, suggesting that fibronectin may have counteracted the hydrophilizing effect of LMWF and improved cell responses. This distinct wettability profile, together with the complementary biological properties of fibronectin and LMWF, may help explain the more favorable endothelial responses observed with their combination.

Surface wettability may also affect liquid spreading during droplet-based cell seeding, potentially influencing cell distribution across stent fragments. This possibility was not directly investigated, as droplet spreading and cell-suspension retention were not quantified. Future studies should assess the contribution of these factors to endothelial colonization, alongside direct characterization of fibronectin organization, potential interactions between fibronectin and LMWF, and the accessibility of cell-adhesive motifs, particularly Arginine-Glycine-Aspartate (RGD) and Proline-Histidine-Serine-Arginine-Asparagine (PHSRN), which contribute to integrin-mediated cell adhesion.

Ibrahim *et al.* reported that optimizing the geometry of laser-induced microgrooves on Co-Cr enhanced endothelial adhesion and increased cell density after 24 h on surfaces with contact angles ranging from 8.9° to 22.2° [38]. Together, these findings illustrate that endothelialization can be enhanced through distinct surface-engineering strategies, with or without substantial changes in wettability. This is consistent with evidence that endothelial responses depend on the interplay between wettability, surface chemistry, functional-group presentation, and surface charge [39–40]. Accordingly, the more hydrophilic LMWF-based coating did not produce the greatest early cell spreading or proliferation, suggesting that wettability alone cannot account for the observed cellular responses.

Surface chemistry may also influence the biological activity of adsorbed fibronectin. Previous studies have shown that surface chemistry can alter fibronectin conformation and integrin-binding specificity [42]. Hao *et al.* further demonstrated that surface-dependent fibronectin adsorption can affect the presentation of integrin-binding motifs, including RGD and PHSRN [31]. These findings offer a potential explanation for why endothelial responses did not follow the wettability ranking of the coatings. The combined coating may create a favorable adhesive interface through effects on surface chemistry and fibronectin presentation. However, because fibronectin conformation and ligand accessibility were not directly assessed in this study, this interpretation remains hypothetical.

The early spreading response is particularly relevant to the establishment of cell-material interactions [43]. Cells on the combined coating occupied a larger projected area at 4 h, whereas the differences between surfaces were no longer significant at 72 h. Thus, the combined interface favored early spreading rather than increasing the final cell area within the observation period. Koenig *et al.* showed that HUVEC attachment and growth on fibronectin-coated polymers correlated with the accessibility of the protein’s cell-binding domain [44]. Their findings suggest that the presentation of fibronectin, rather than its presence alone, can influence endothelial responses. In our system, the greater early spreading on fibronectin-LMWF is consistent with a contribution from adhesive protein-cell interactions [45]. However, the absence of a fibronectin-only condition in the spreading analysis prevents us from separating this contribution from that of LMWF.

Overall, these findings identify fibronectin precoating as a strategy to enhance LMWF retention on Co-Cr stents and improve endothelial adhesion, spreading and lamellipodia formation while largely preserving the wettability of the underlying substrate. Given the importance of timely re-endothelialization for vascular healing after stent implantation [2], this approach warrants further investigation. Future studies should clarify the mechanisms underlying these responses and determine whether the coating influences smooth muscle cell behavior and neointimal hyperplasia.

### ❖ The combined coating benefits distinct stages of endothelial colonization

Endothelial colonization followed a time-dependent pattern across all surface conditions, with cell density decreasing between 4 and 48 h before recovering at 72 h. This initial decline may reflect cell loss or adaptation to the surface, consistent with previous reports of early endothelial detachment from metallic endovascular implants [45]. Notably, the fibronectin-LMWF coating maintained higher cell densities than the LMWF coating alone at 24 and 48 h. The subsequent recovery in cell density was followed by an increased EdU incorporation between 72 and 96 h, which was greater with the combined coating than with either component alone, indicating enhanced proliferative activity at later time points. Moreover, LDH release was lower from fibronectin-LMWF-coated than from fibronectin-coated stents at 72h, suggesting limited membrane damage under the tested conditions. Collectively, these findings highlight the potential of fibronectin-LMWF surface modification to support endothelial colonization over time, combining higher early cell density with enhanced later proliferative activity.

These findings extend our previous observations that soluble LMWF promotes HUVEC proliferation, spreading, and migration [14] to a surface-functionalization strategy. However, the effects of soluble fucoidan depend on the growth stimulus enhancing serum and FGF-1-induced growth while inhibiting FGF-2-induced proliferation [46]. This context dependence is relevant to the present surface-bound formulation, in which fibronectin precoating was associated with greater EdU incorporation than LMWF alone. Differences in LMWF loading or surface presentation may contribute to this response; however, the present data do not distinguish between increased LMWF availability and effects associated with fibronectin-mediated cell adhesion.

MMP measurements provided further insight into the remodeling-associated response to surface functionalization [47]. The fold change in MMP-2 concentration (48 h/24 h) was higher in the fibronectin-LMWF-coated group, whereas MMP-9 concentration fold changes did not differ significantly across coating conditions. This profile is consistent with our previous findings that soluble LMWF enhanced HUVEC migration and increased MMP-2 expression and secretion, while MMP-2 knockdown attenuated migration [14]. Bouvard *et al.* further implicated PI3K/AKT signaling in LMWF-induced endothelial migration [48]. These findings provide a mechanistic context for the responses observed here; however, the temporal increase in MMP-2 did not translate into a significant difference relative to the LMWF-coated group, and MMP concentrations do not establish enzymatic activity. Thus, the present data identify a remodeling-associated response but do not demonstrate enhanced endothelial motility.

Because MMPs have been implicated in SDC ectodomain shedding, we performed dot blot analyses to assess soluble SDC-1 and SDC-4 levels across coating conditions. In particular, MMP-9 has been shown to mediate SDC-4 shedding in endothelial cells, contributing to alterations in the endothelial glycocalyx [47]. In our previous study, soluble LMWF increased soluble SDC-1 levels while decreasing soluble SDC-4 levels in HUVEC-conditioned media [16]. However, the present analyses revealed no significant differences in SDC-1 or SDC-4 shedding among the coating conditions. This discrepancy may reflect differences in LMWF presentation, cellular exposure, or culture conditions, as surface-immobilized and soluble LMWF may elicit distinct cellular responses. Nevertheless, the absence of detectable differences in soluble SDC levels does not exclude changes in membrane-associated SDC expression or shedding activity. Further studies are needed to clarify how LMWF presentation influences SDC regulation and whether this process contributes to endothelial responses to the coated stents.

### ❖ Coating-associated changes in endothelial organization

Confocal microscopy revealed coating-associated differences in VE-cadherin localization at cell-cell contacts, F-actin distribution, and integrin α6/β1 and SDC-1/SDC-4 staining patterns, highlighting variations in endothelial junctional organization, cytoskeletal architecture, and syndecan distribution. These observations are consistent with the broader understanding that cellular responses to biomaterials depend on the spatial organization of adhesive interactions [50]. Using a non-endothelial cell model with a fixed fibronectin surface density, Keselowsky *et al*. showed that surface chemistry influences integrin binding, focal adhesion protein recruitment, and FAK signaling [50]. Their findings suggest that the biological presentation of adhesive ligands can affect adhesion complex assembly independently of surface density, providing a mechanistic framework for our observations, although these processes were not directly investigated in our model.

VE-cadherin staining appeared more intense on LMWF-coated and fibronectin-LMWF-coated surfaces, with a thicker pattern in the combined coating condition, potentially indicating differences in junctional organization. F-actin fibers were relatively linear and homogeneously distributed throughout the cytoplasm on uncoated Co-Cr and fibronectin-coated surfaces, whereas LMWF-coated and fibronectin-LMWF-coated exhibited more pronounced submembranous localization. These patterns may reflect coating-dependent differences in cytoskeletal organization and cell-cell contacts, although their functional significance remains to be established. Integrin β1 and α6 staining showed more peripheral, membrane-associated localization on coated surfaces, particularly with the combined coating, whereas LMWF-coated groups exhibited a more diffuse intracellular pattern. Notably, these qualitative differences were not accompanied by significant changes in total VE-cadherin or integrin β1/α6 abundance.

Similarly, Rohringer *et al.* reported improved HUVEC adhesion on chorion-derived extracellular matrix-coated expanded polytetrafluoroethylene (ePTFE) grafts without significant increases in integrin α5, β1, or β3 levels [51]. Fucoidan-mediated effects on endothelial adhesion and integrin regulation have also been reported [37,52]. However, whether the patterns observed here reflect changes in protein organization or function remains unclear.

The phospho-paxillin/total paxillin ratio did not differ significantly between conditions, providing no evidence that decreased paxillin phosphorylation underlies the observed cellular responses. Crawford *et al.* reported that paxillin and focal adhesion kinase (FAK) can localize at cadherin-containing cell-cell junctions, suggesting a potential role in coordinating cell-matrix and cell-cell adhesion [53]. Accordingly, the junction-associated VE-cadherin staining observed here may reflect differences in cell-cell contact organization and potentially involve crosstalk with focal adhesion-associated proteins. However, the absence of significant changes in paxillin phosphorylation and direct evidence of paxillin-VE-cadherin interactions precludes establishing this mechanism. Moreover, VE-cadherin localization at cell-cell contacts does not, by itself, demonstrate improved endothelial barrier function [54].

The SDC findings provide a potential link between the material interface and endothelial signaling. LMWF has previously been implicated in endothelial migration and angiogenesis through pathways involving heparanase and SDC-4. We previously reported that soluble LMWF increased SDC-1 expression and ectodomain shedding in HUVECs, while decreasing SDC-4 levels [16], suggesting differential regulation of these proteoglycans. In the present study, cells on the combined coating exhibited broader intracellular staining for SDC-1 and SDC-4. However, these qualitative patterns reflect protein distribution and cannot be interpreted as changes in total expression or ectodomain shedding. Consistent with this distinction, soluble SDC-1 and SDC-4 levels did not differ significantly among coating conditions. Thus, the observed staining patterns identify syndecan localization as a potential avenue for further investigation but do not establish changes in syndecan expression or shedding.

### ❖ Endothelial mediator profiles and SMC responses to conditioned media

To assess the paracrine effects of the endothelial layer, we examined the soluble mediator profiles in the HUVEC-conditioned media and then their effects on smooth muscle cell (SMC) migration and metabolic activity. VEGF-A secretion was transiently increased at 48 h on uncoated stents, which exhibited lower endothelial cytocompatibility than the coated surfaces. This observation raises the hypothesis that VEGF-A release may reflect a stress-associated endothelial response, potentially linked to its established roles in angiogenesis and tissue repair [55]. However, this interpretation remains speculative and requires further investigation. The fibronectin-LMWF coating did not significantly increase the levels of the inflammatory mediators (MCP-1/CCL2, RANTES/CCL5, IL-6, TNF-α, or IL-10) compared with the other surface conditions. Although IL-8/CXCL8 levels increased over time in this group, similar changes occurred on uncoated and LMWF-coated stents, indicating that this response was not specific to the combined coating. IL-8/CXCL8 has been shown to promote endothelial migration and cytoskeletal reorganization through PI3K-Rac1/RhoA signaling [56]. In the present study, lamellipodia formation increased up to 72 h of endothelial colonization on fibronectin-LMWF-coated stents, while F-actin showed a more pronounced submembranous localization on both LMWF-and fibronectin-LMWF-coated stents. These morphological observations coincided with temporal changes in IL-8/CXCL8 levels and may suggest an association between cytokine secretion and endothelial cell remodeling. However, the absence of a coating-specific IL-8/CXCL8 response and the lack of direct functional evidence preclude establishing a causal relationship.

Finally, conditioned-medium experiments assessed whether coating-associated endothelial responses influenced SMC behavior. No significant differences in SMC migration or metabolic activity were detected among the four surface conditions, indicating that the endothelial responses associated with fibronectin-LMWF were not accompanied by detectable changes in these SMC endpoints. In our previous study, direct exposure to soluble LMWF reduced vascular SMC migration and MMP-2 expression while promoting endothelial migration [14]. In the present study, however, SMCs were exposed to endothelial-conditioned media containing cell-derived factors and potentially material released from the coatings. Differences in effective LMWF exposure and medium composition may therefore help explain the absence of a detectable inhibitory effect. Accordingly, fibronectin-LMWF surface modification did not produce detectable additional effects on SMC migration or metabolic activity under the conditions tested. Whether this coating can limit neointimal growth remains to be investigated in an appropriate vascular model.

## 5. Conclusion

Sequential biomimetic fibronectin-LMWF coating enhanced LMWF deposition and retention on Co-Cr stent surfaces, while promoting endothelial early cell spreading, lamellipodia formation, and subsequent proliferative activity. These benefits were accompanied by changes in MMP-2 and IL-8/CXCL8 levels and in the organization and localization of cytoskeletal, junctional, and adhesion-associated proteins, without a coating-specific increase in the measured inflammatory mediators. Importantly, endothelial-conditioned media from fibronectin-LMWF-coated stents did not induce greater SMC migration than media from the other conditions. Collectively, these findings establish fibronectin-assisted LMWF deposition as a promising strategy to enhance endothelial colonization of metallic stents while preserving a favorable inflammatory profile. Under physiological flow and in vascular implantation models are required to establish coating stability, hemocompatibility, and its effects on vascular healing and neointimal hyperplasia. Further studies under physiological flow and in vascular implantation models are required to establish coating stability, hemocompatibility, and its effects on vascular healing and neointimal hyperplasia.

**************************************************************************

## Research Highlights

### ❖ What is the current knowledge?

● LMWF promotes endothelial cell spreading, proliferation and migration while inhibiting SMC migration in vitro.
● Fucoidan binds to fibronectin at multiple sites.
● Fucoidan-polydopamine coating on Mg-Zn-Y-Nd alloy enhances endothelialization and may limit neointimal hyperplasia.

### ❖ What is new here?

● Fibronectin precoating enhances LMWF immobilization and retention on Co-Cr stents.
● Fibronectin-LMWF coating promotes endothelial colonization and proliferation with high cytocompatibility.
● Fibronectin-LMWF coating promotes HUVEC spreading, lamellipodia formation, and F-actin remodeling.
● Fibronectin-LMWF coating modulates the endothelial secretome, including IL-8/CXCL8 and MMP-2.

**************************************************************************

## Acknowledgements

This work was supported by the Direction de la Recherche et des Études Doctorales (Ministère de l’Enseignement Supérieur et de la Recherche), Université Sorbonne Paris Nord (USPN), and Institut National de la Santé et de la Recherche Médicale (INSERM). Additional financial support was provided by the Institut Universitaire de France (IUF) and the Fondation de l’Avenir. M. Saadi received fellowships from the ministère de l’Enseignement Supérieur et de la Recherche and USPN (SREI 2023). C. Boisson-Vidal’s salary was funded by the Centre National de la Recherche Scientifique (CNRS).

The authors thank AndraTec Medical Devices and Prof. Valérie Besnard (INSERM UMRS 1272) for their valuable collaboration. We also thank Dr. Michael Trichet (Institut de Biologie Paris Seine, Sorbonne University) for his assistance with electron microscopy, with additional support from the Île-de-France Region and the CNRS. The authors are grateful to François Lanthiez, head of the IMA’CRI imaging platform (INSERM UMR 1149), for his assistance with confocal microscopy. Finally, we sincerely thank Prof. Carole Planes, director of the Hypoxie & Poumon laboratory (INSERM UMRS 1272), for providing access to research facilities.

## CRediT authorship contribution statement

Mehdi SAADI: Writing – original draft, Visualization, Resources, Methodology, Investigation, Formal analysis, Data curation. Raymond HADDAD: Validation, Investigation, Formal analysis, Roles/Writing – original draft. Nour ZEAITER: Investigation. Oualid HADDAD: Investigation, Methodology, Roles/Writing – original draft, Data curation. Divya PRABHAKAR REDDY: Investigation, Roles/Writing – original draft, Software, Data curation. Jean Sébastien BAUMANN: Investigation, Methodology, Data curation. Fahima IRKI: Investigation. Jessica BAMBA-FUNCK: Formal analysis, Investigation. Angela SUTTON: Resources, Methodology, Roles/Writing – original draft. Céline FALENTIN-DAUDRE: Methodology, Roles/Writing – original draft. Kévin BASSAND: Investigation, Methodology, Validation, Data curation, Roles/Writing – original draft, Writing – review & editing. Catherine BOISSON VIDAL: Methodology, Validation, Roles/Writing – original draft, Writing – review & editing. Hanna HLAWATY: Conceptualization, Roles/Writing – original draft, Writing – review & editing, Supervision, Validation, Project administration, Investigation, Funding acquisition, Formal analysis.

## Data Availability Statement

The data supporting the findings of this study are available from the corresponding author upon reasonable request.

---

**Supplementary figure 5. Time-dependent endothelial colonization of uncoated and coated Co-Cr stents.**

**(A)** Representative fluorescence microscopy (FM) images showing HUVEC distribution on uncoated, fibronectin-coated, LMWF-coated, and fibronectin-LMWF-coated Co-Cr stents at 4, 48, and 72 h after seeding. HUVECs were labelled with CellTracker Green, and white dashed lines delineate the stent boundaries. Scale bars: 100 µm.

**Supplementary figure 6. Endothelial adhesion-associated protein levels and immunofluorescence isotype controls.**

**(A-C)** Representative Western blotting images and densitometric quantification of VE-Cadherin **(A)**, integrin β1 **(B)**, and integrin α6 **(C)** protein levels in HUVECs cultured on uncoated, fibronectin-coated, LMWF-coated, and fibronectin-LMWF-coated Co-Cr stents for 48 h. Protein levels were normalized to actin then to the uncoated condition and are expressed as arbitrary units (a.u.). Data are mean ± SEM (n ≥ 3). One-way ANOVA with Tukey’s multiple-comparisons test. ns, not significant. **(D)** Representative Western blotting images and densitometric quantification of phosphorylated paxillin (P-Paxillin) relative to total paxillin in HUVECs cultured on uncoated, fibronectin-coated, LMWF-coated, and fibronectin-LMWF-coated Co-Cr stents for 48 h. P-Paxillin levels were normalized to total paxillin, corrected for actin loading, and subsequently normalized to the uncoated condition. Data are expressed as arbitrary units (a.u.) and presented as mean ± SEM (n = 4). One-way ANOVA with Tukey’s multiple-comparisons test. ns, not significant. **(E)** Representative fluorescence microscopy (FM) images of HUVECs incubated with IgG2a_k_ and IgG1_k_ isotype controls. Nuclei were counterstained with DAPI (blue), and white dashed lines delineate the stent boundaries. Scale bars: 100 µm.

**Supplementary figure 7. Growth factor, chemokine, and cytokine concentrations in endothelial culture supernatants.**

Quantification of vascular endothelial growth factor A (VEGF-A) **(A)**, fibroblast growth factor-2 (FGF-2) **(B)**, monocyte chemoattractant protein-1 (MCP-1/CCL2) **(C)**, regulated upon activation, normal T cell expressed and secreted (RANTES/CCL5) **(D)**, interleukin-8 (IL-8/CXCL8) **(E)**, interleukin-6 (IL-6) **(F)**, tumor necrosis factor-α (TNF-α) **(G)**, and interleukin-10 (IL-10) **(H)**, in culture supernatants from HUVECs cultured on uncoated, fibronectin-coated, LMWF-coated, and fibronectin-LMWF-coated Co-Cr stents. HUVECs were initially cultured for 24 h in complete medium, followed by 24 and 48 h in serum-free medium before conditioned-medium collection. Analyte concentrations were measured using the Meso Scale Discovery (MSD) multiplex immunoassay and are expressed as pg/mL. Data are mean ± SEM (n = 3). Two-way ANOVA with Tukey’s multiple-comparisons test. \**p* < 0.05; \*\**p* < 0.01; ns, not significant.

