## Supplementary figures and images for "Fibronectin-Assisted Coating of Low-Molecular-Weight Fucoidan Creates a Biomimetic Interface for Endothelialization of Cobalt-Chromium Stents"

### Supplemental Figure 5

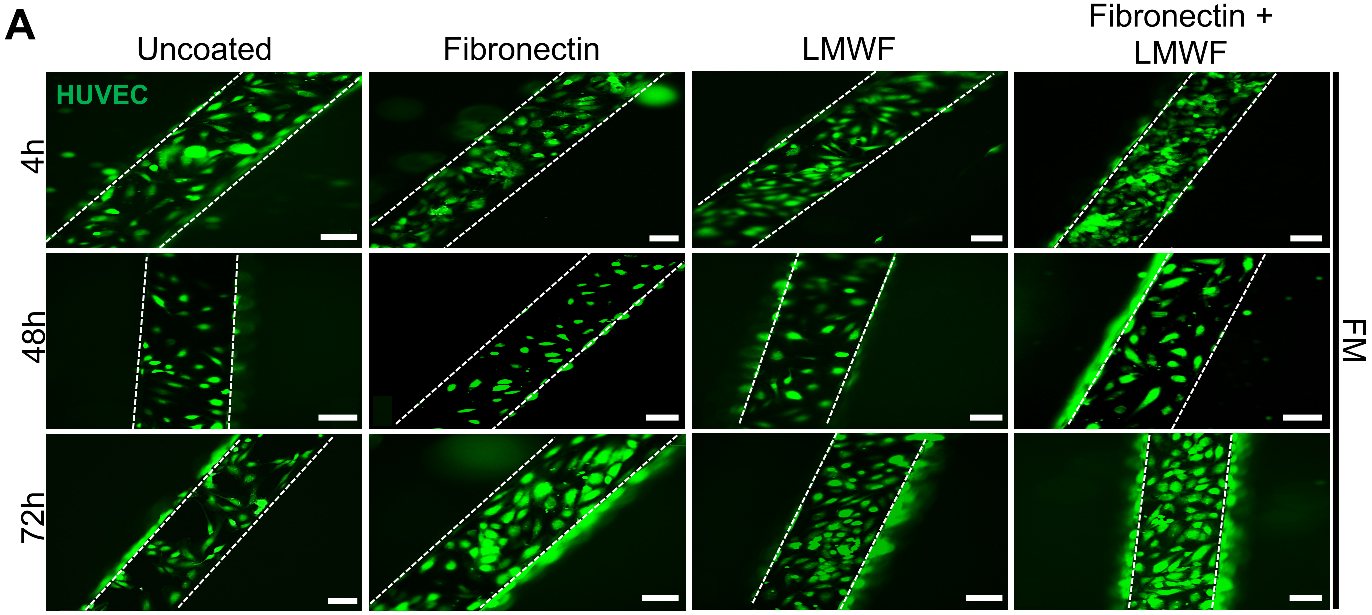

A
Uncoated
Fibronectin
LMWF
Fibronectin +
LMWF
HUVEC
4h
48h
72h
FM
