## Supplemental Figure 6 for "Fibronectin-Assisted Coating of Low-Molecular-Weight Fucoidan Creates a Biomimetic Interface for Endothelialization of Cobalt-Chromium Stents"

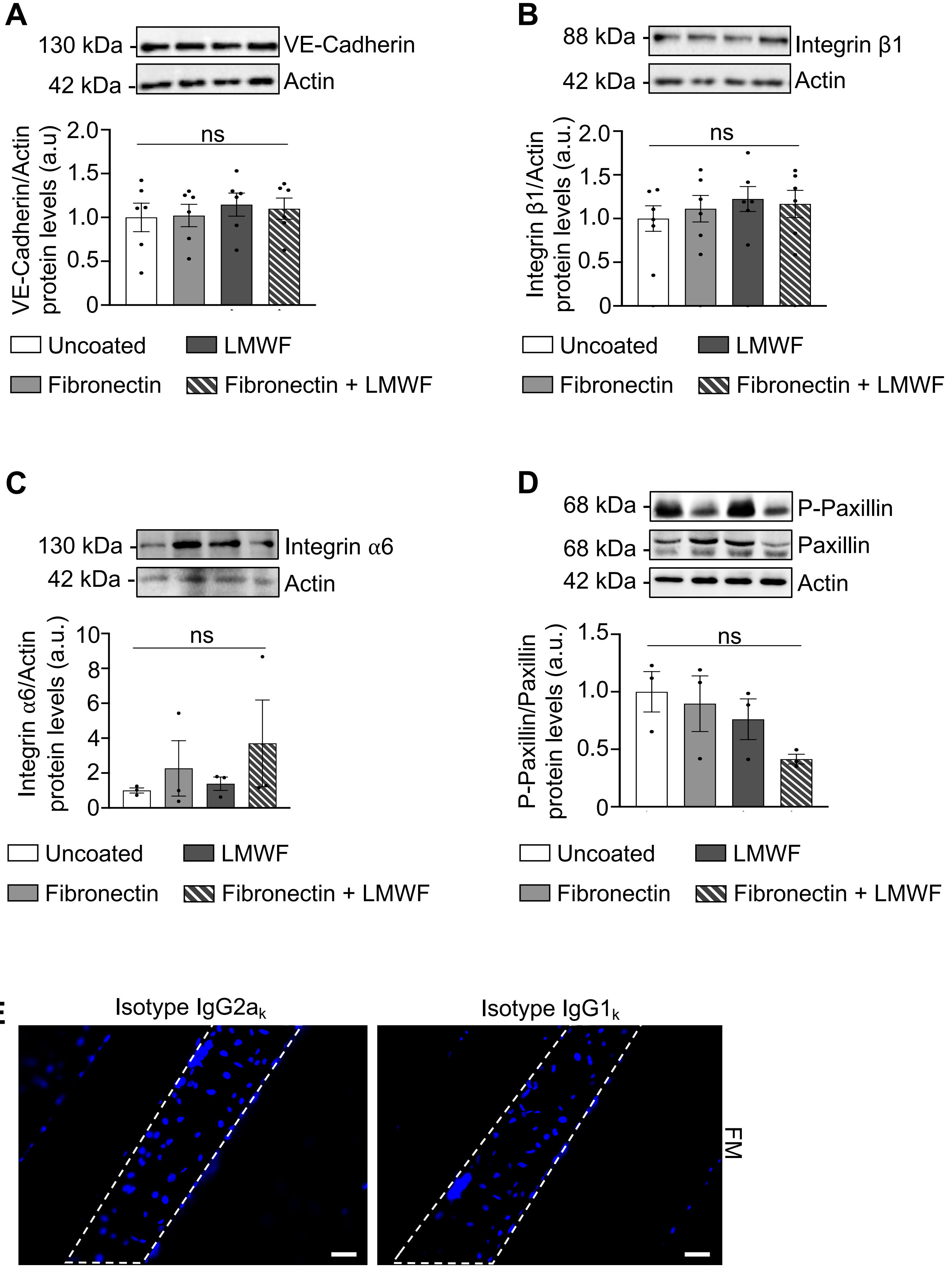
A
130 kDa
VE-Cadherin
42 kDa
Actin
VE-Cadherin/Actin protein levels (a.u)
ns
B
88 kDa
Integrin β1
42 kDa
Actin
Integrin β1/Actin protein levels (a.u.)
ns
Uncoated
LMWF
Fibronectin
Fibronectin + LMWF
C
130 kDa
Integrin α6
42 kDa
Actin
Integrin α6/Actin protein levels (a.u.)
ns
D
68 kDa
P-Paxillin
68 kDa
Paxillin
42 kDa
Actin
P-Paxillin/Paxillin protein levels (a.u.)
ns
E
Isotype IgG2ak
Isotype IgG1k
FM
