## Supplemental Figure 7 for "Fibronectin-Assisted Coating of Low-Molecular-Weight Fucoidan Creates a Biomimetic Interface for Endothelialization of Cobalt-Chromium Stents"

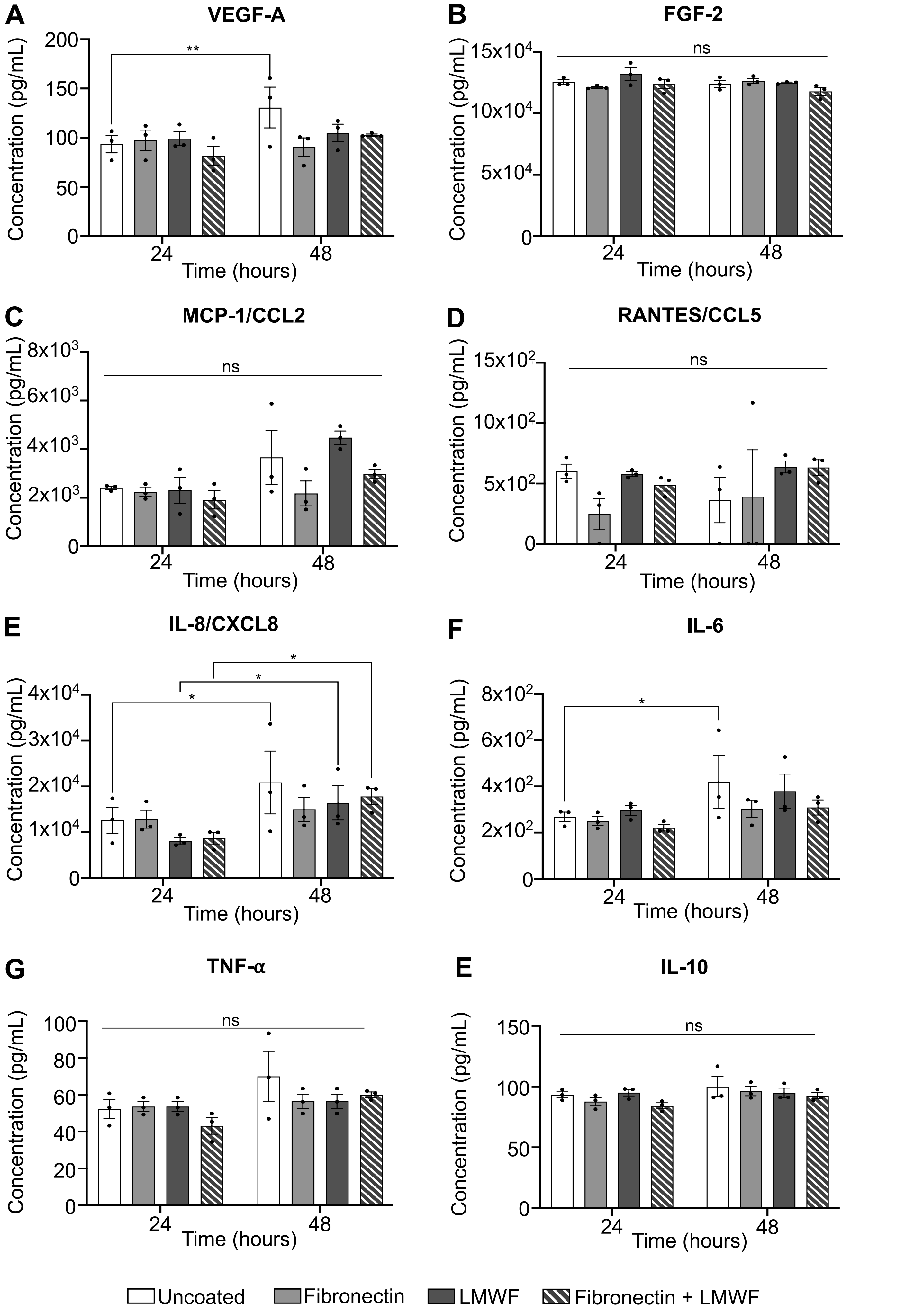
A
VEGF-A
**
Concentration (pg/mL)
Time (hours)
B
FGF-2
ns
C
MCP-1/CCL2
ns
D
RANTES/CCL5
ns
E
IL-8/CXCL8
*
F
IL-6
*
G
TNF-α
ns
E
IL-10
ns
Uncoated
Fibronectin
LMWF
Fibronectin + LMWF
